# Mechanistic basis for the spatiotemporal assembly of the Bloom Syndrome complex on ultrafine DNA bridges

**DOI:** 10.64898/2026.09.10.750333

**Authors:** Umit Aliyaskarova, Özgün Özer, Mahafuzul Sumon, Eleftheria Karanika, Adrijana Crncec, Zuzanna Kozik, Theo Roumeliotis, Jyoti Choudhary, María Fernández-Casañas, Matthew Day, Antony W Oliver, Jonathon Pines, Kok-Lung Chan

## Abstract

The BTRR (BLM/TOP3A/RMI1/RMI2) complex resolves DNA replication and recombination intermediates, as well as mitotic DNA intertwinements, yet how its diverse activities are regulated remains unclear. Here, we identify key determinants of stable assembly of the BTRR complex on ultrafine DNA bridges (UFBs). We reveal that the RMI1-RMI2 subcomplex acts as a molecular reader of hydrophobic-rich short linear motifs (SLiMs) embedded within PICH and FANCM, thereby providing a conserved mechanism for engaging distinct Superfamily2 DNA translocases during chromosome segregation and DNA repair. During UFB resolution, SLiM binding stabilises the PICH-BTRR complex assembly and supports essential interactions among BLM, PICH, and DNA. Persistent PICH-BTRR complex association post-mitosis would compromise interphase DNA damage responses and is prevented by active PICH nuclear exclusion. Our findings define the molecular basis of UFB-binding complex assembly and highlight its spatiotemporal control required to specialise BTRR complex functions.

## Introduction

The Bloom syndrome helicase BLM, together with Topoisomerase 3A, RMI1, and RMI2, collectively known as the BTRR complex^1^, plays multifaceted roles in maintaining chromosome stability across both interphase and mitosis. Its well-established functions include replication fork recovery, suppression of hyper-recombination, and ultrafine DNA bridge (UFB) resolution^2–12^. Defects in any of these pathways can severely threaten genomic integrity, driving the growth defects and cancer predisposition characteristic of Bloom syndrome^13–15^. Unsurprisingly, these diverse roles of the BTRR complex depend on chromatin context, cell cycle stage and specific protein-protein interactions (PPIs) made with different binding partners.

During DNA replication, many key repair factors involved in DNA end-resection and homologous recombination, such as EXO1, MRN, TOPBP1, and RAD51, have been found to physically and functionally interact with BLM. These PPIs are primarily mediated by distinct sites in the N-terminal unstructured region of BLM^16–21^. In addition, the BTRR complex can bind to the Superfamily 2 (SF2) DNA helicase FANCM via the RMI1-RMI2 subcomplex for damaged fork repair^22,23^. During mitosis, particularly when cells undergo sister chromatid disjunction, the BTRR complex interacts with the SNF2-like DNA translocase PICH (also known as ERCC6L) together with Polo-like kinase 1 (PLK1) and RIF1-PP1, forming a mitosis-specific complex known as ‘the ultrafine DNA bridge (UFB)-binding complex’^2,24,25^. The UFB-binding complex is a potent molecular machine capable of resolving interlinked sister DNA molecules within minutes^4^. However, its premature activation prior to anaphase onset causes severe centromere disintegration^26^. We and others have shown that this mitotic catastrophe is suppressed by CDK1- and PLK1-mediated hyperphosphorylation, which inhibits PPIs among PICH, the BTRR complex, and the RIF1-PP1 protein^25–28^. Thus, while strict spatiotemporal control of the UFB-binding complex assembly is vital for mitosis, how cells govern the engagement of the BTRR complex and RIF1-PP1 with PICH on anaphase bridges remains poorly understood.

Both PICH and BLM belong to the SF2-helicase superfamily; however, PICH lacks canonical DNA unwinding activity^29^. Instead, it exhibits functions in nucleosome remodelling, triplex DNA displacement and Holliday junction branch migration^29–31^. Despite being a chromatin remodeler, PICH is highly unusual among the known SNF2-like translocases in mammalian cells in that it predominantly localises outside the nucleus throughout interphase^24,32^. The biological significance of this nuclear exclusion remains elusive. Structurally, the core catalytic domains of PICH reside in its N-terminus, whereas its C-terminus is predicted to be largely disordered and lacks well-defined functions^33^ (Supplementary Fig.1a). Our previous study identified a conserved short linear motif (SLiM) within the C-terminally disordered region of PICH that conforms to the consensus of the ABBA motif found in Cyclin A, BUBR1, BUB1 and Acm1^34,35^ (Fig. 1a). Since ABBA motifs bind and regulate CDC20, an activator of the Anaphase Promoting Complex/Cyclosome (APC/C)^34,35^, this raises the possibility that PICH may also be involved in the Spindle Assembly Checkpoint (SAC)^24^, but compelling evidence supporting this proposal is lacking^36^. On the other hand, a recent report proposed that the region harbouring the ABBA motif in PICH is important for interactions with BLM and RIF1^37^.

**Figure 1.**
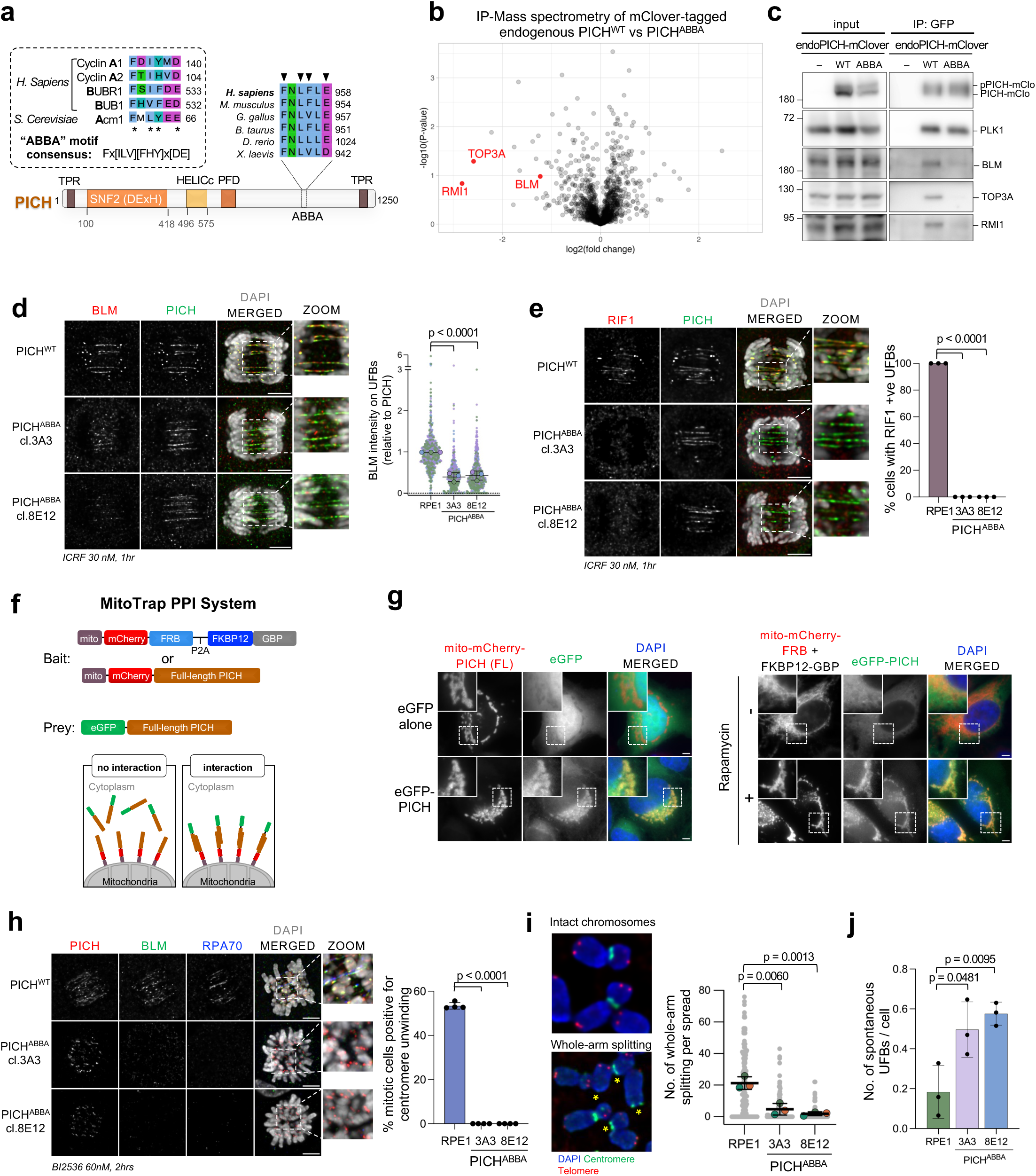
Activation of the UFB-binding complex requires the PICH ABBA motif. **a.** Sequence alignments of the PICH ABBA motifs across species. Arrowheads indicate the conserved residues Phe953, Leu955, Phe956, and Glu958 of PICH found in ABBA consensus sequences (inset box). Domain organisation of PICH is shown (TPR: tetratricopeptide repeats; SNF2: sucrose non-fermenting family domain; PFD: PICH family domain; HELICc: helicase superfamily C-terminal domain). Inset box: consensus sequences of ABBA motifs found in Cyclin A, BUBR1, BUB1, and Acm1. **b.** Volcano plot of a mass spectrometry-based proteomic comparison of Nocodazole-arrested mitotic extracts PICH wild-type (PICH^WT^) and ABBA mutant (PICH^ABBA^) cells after immunoprecipitation of endogenously mClover3-tagged PICH proteins. The full list of pulled-down proteins is provided in Supplementary Data 2. **c.** Co-immunoprecipitation of endogenous mClover3-tagged PICH in nocodazole-arrested mitotic RPE1 cells treated with 100nM of PLK1i (BI2536) for the last 1hr. **d.** UFB localisation of BLM and PICH in PICH^WT^ and PICH^ABBA^ RPE1 anaphase cells after ICRF193 treatments. Left: representative images of UFB staining. Right: relative BLM intensity on UFBs (normalised to the average BLM intensity in PICH^WT^ RPE1 cells). Two PICH^ABBA^ mutant clones were measured. Total number (n) of UFBs measured across three independent experiments: PICH^WT^, n = 618, 81 cells; PICH^ABBA^ 3A3, n = 534, 68 cells; PICH^ABBA^ 8E12, n = 628, 74 cells; mean ± S.D. is shown. **e.** UFB localisation of RIF1 in PICH^WT^ and PICH^ABBA^ RPE1 anaphase cells after ICRF193 treatments. Left: representative images of UFB staining. Right: percentage of anaphase cells showing ≥ 1 RIF1-positive UFB. Total number (n) of cells analysed across three independent experiments: PICH^WT^ n = 96; PICH^ABBA^ 3A3, n = 76; PICH^ABBA^ 8E12, n = 70; mean ± S.D. is shown. **f.** Schematic diagram showing the MitoTrap Protein-Protein Interaction (PPI) System. eGFP-PICH (prey), is co-expressed with the indicated bait proteins. As a control, Mito-mCherry-FRB is immobilised on mitochondria, whereas FKBP12-GBP(GFP-binding protein) is used to capture eGFP-PICH. In the presence of rapamycin, the FKBP12-GBP complex captures eGFP-PICH, which localises to mitochondria via FRB-FKBP12 dimerisation. If PICH forms homodimers or multimers, eGFP-PICH will also be immobilised to mitochondria when co-expressed with Mito-mCherry-PICH. **g.** Representative examples of HeLa cells expressing eGFP-PICH and mito-mCherry fused with either FRB-P2A-FKBP12-GBP or full-length PICH. Left: eGFP-PICH, but not eGFP alone, localises to mitochondria after mito-mCherry-PICH expression. Right: rapamycin-induced FRB-FKBP12 dimerisation triggers mitochondrial localisation of eGFP-PICH captured by the FKBP12-GBP bait protein. **h.** Defective K-chromatin localisation and centromere unwinding induced by PLK1 inhibition in PICH^ABBA^ cells. Left: Representative images of BLM and RPA70 at centromeres/centromeric threads in PICH^WT^ and PICH^ABBA^ RPE1 cells after BI2536 treatments. Right: Percentages of mitotic cells exhibiting centromeric unwinding. Total number (n) of cells analysed across four independent experiments: PICH^WT^, n = 241; PICH^ABBA^ 3A3, n = 165; PICH^ABBA^ 8E12, n = 155; mean ± S.D. is shown. **i.** Defective whole-chromosome arm splitting induced by PLK1 inhibition. Left: representative images of intact and whole-arm-split chromosomes (asterisks). Right: quantification. Total number (n) of spreads analysed across three independent experiments: PICH^WT^, n = 155; PICH^ABBA^ 3A3, n = 103; PICH^ABBA^ 8E12, n = 136; mean ± S.D. is shown. **j.** Number of spontaneous UFBs in PICH^WT^ and PICH^ABBA^ cells. Total number (n) of anaphase cells scored across 3 independent experiments: PICH^WT^, n = 182; PICH^ABBA^ 3A3, n = 175; PICH^ABBA^ 8E12, n = 196; mean ± S.D. is shown. All p-values are calculated by unpaired two-tailed t-test; Scale bars, 5 μm.

Here, we have investigated the molecular functions of the PICH ABBA motif and the mechanism driving UFB-binding complex assembly. By combining structural modelling, sequence conservation analysis, and cellular protein-protein interaction assays, we define the critical determinants of UFB-binding complex assembly in human mitotic cells. We identify the RMI1-RMI2 heterodimer as a specialised molecular reader for sequence-unrelated, hydrophobic-rich SLiMs found in both FANCM and PICH SF2 DNA translocases. This essential SLiM-recognition module enables the BTRR complex to alternate between partners, executing distinct activities in interphase and mitosis, respectively. Beyond providing a new structural blueprint for the UFB-binding complex, our findings underscore the importance of the spatiotemporal exchange between different DNA translocases for genome protection.

## Results

### Homozygous mutations of the PICH ABBA motif do not compromise the SAC

To elucidate the cellular functions of the PICH ABBA motif, we generated knock-in mutations in both alleles of PICH in RPE1 FRT/TR cells using CRISPR-mediated genome editing (Supplementary Fig. 1b). Four conserved residues in the ABBA motif (Phe953, Leu955, Phe956, and Glu958) were substituted with alanine, hereafter referred to as the PICH^ABBA^ mutant (Fig. 1a). Whole-exon sequencing of the entire PICH gene confirmed that only the four amino acids had been mutated (Supplementary Fig. 1c and Supplementary Data 1). We observed slow-growth phenotypes and reduced colony formation efficiencies in both PICH^ABBA^ mutant clones (Supplementary Figs. 1d, 1e), but there were no obvious changes in the cell cycle distribution (Supplementary Fig. 1f). We tested whether mutating PICH had affected the spindle assembly checkpoint (SAC) by assaying the ability of cells to delay in mitosis when treated with nocodazole but found no detectable defects (Supplementary Fig. 1f). Moreover, mass spectrometry analysis of immunoprecipitated endogenous PICH^WT^ did not detect any CDC20 peptides (see below; Supplementary Data 2), nor did reciprocal mitotic CDC20 immunoprecipitants contain PICH peptides (Supplementary Data 3). Altogether, we conclude that the ABBA motif of PICH is unlikely to function for CDC20 binding and SAC regulation.

### The assembly and activation of UFB-binding complexes require the ABBA motif

To identify what proteins did bind the PICH ABBA motif, we analysed immunoprecipitants of endogenous PICH proteins pulled down from PICH^WT^ and PICH^ABBA^ RPE1 mitotic cells by mass spectrometry (Supplementary data 2). An mClover3-tag was inserted at the C-terminus of PICH in both cell lines to enhance the efficiency and specificity of immunoprecipitation (Supplementary Fig. 2a). The mClover3 tag did not alter the normal localisation of PICH in either interphase or mitotic cells (Supplementary Fig. 2b). Mass spectrometry analysis revealed that the PICH^ABBA^ mutant was perturbed in interacting with components of the BTRR complex, including BLM, TOP3A, and RMI1 (Fig. 1b). Co-immunoprecipitation experiments further confirmed the reduced interaction with BTRR complex subunits under conditions of PLK1 inhibition (Fig. 1c), which have shown to promote the interaction between PICH and the BTRR complex^27^. Consistent with these findings, the localisation of BLM protein to anaphase UFBs was dramatically reduced in PICH^ABBA^ cells (Fig. 1d and Supplementary Fig. 2c). A slight reduction in UFB binding by the PICH^ABBA^ mutant protein was observed, but less pronounced than that observed for BLM (Supplementary Fig. 2c). It is worth noting that the binding of BLM to UFBs was not fully abolished (Fig. 1d), whereas the recruitment of RIF1 was completely ablated (Fig. 1e). This indicated that there may be other PICH-binding interfaces in the BTRR complex. Given that BLM and RIF1 localisations to UFBs are independent of each other^25^ but both of their recruitments are impaired in PICH^ABBA^ mutant cells, these findings imply that BLM and RIF1 likely interact with PICH in a mutually exclusive manner. Immunofluorescence imaging reveals a very similar localisation pattern for both BLM and RIF1 along UFB^25^, which could be due to stochastic interactions with PICH. Alternatively, we speculated that this could result from dimerisation or oligomerisation of PICH, thereby enabling the co-recruitment of these proteins to UFBs. To test the potential of PICH to dimerise in cells, we used a modified MitoTrap system^38^. Briefly, ectopically expressed PICH proteins were tagged with either eGFP or with Tom70p-mCherry (hereafter referred to as Mito-mCherry) that can immobilise proteins of interest onto mitochondrial surfaces (Fig. 1f). We found that eGFP-PICH, but not eGFP proteins, localised to the mitochondria when Mito-mCherry-PICH was co-expressed, indicating that PICH homodimerisation can occur (Fig. 1g; left panels). As a control, eGFP-PICH alone did not bind to mitochondria but translocated there in the presence of Mito-mCherry-FRB and FKBP12-GBP (<u>G</u>FP-<u>b</u>inding <u>p</u>rotein) when rapamycin was added to induce FRB-FKBP12 dimerization (Figs. 1f, 1g; right panels). This result shows that PICH can form dimers or oligomers in cells and could support the co-recruitment of both BLM/BTRR complexes and RIF1 on ultrafine DNA bridges.

Next, we examined the functionality of the PICH^ABBA^ mutant. Previously, we showed that UFB-binding complexes are prematurely activated by PLK1 inhibition, leading to centromere disintegration in mtiosis^26^. We adopted this mitotic catastrophe response as a functional assay for PICH. When we inhibited PLK1, unlike the wildtype cells, PICH^ABBA^ mutants displayed very low frequencies of centromeric thread formation and whole-chromosome arm splitting - markers of centromere disintegration^26^(Figs. 1h, 1i). Moreover, persistent UFB formations were detected in PICH^ABBA^ cells (Fig. 1j). These results indicate that the PICH^ABBA^ mutant was barely functional and point to defects in the overall activity of the UFB-binding complex. We conclude that the PICH ABBA motif is crucial for the assembly and activation of the UFB-binding complex.

### AlphaFold predicts RMI1-RMI2 as a SLiM reader

To gain insights into the underlying mechanism of UFB-binding complex formation, we employed AlphaFold (AF) to generate structural predictions. Here, we focused on the interaction between the BTRR complex and PICH because both the BLM and TOP3A subunits are believed to play important catalytic roles in UFB resolution^2,4,26,27^. Five structural models comprising full-length BLM, TOP3A, RMI1, RMI2 and PICH were generated using AlphaFold-Multimer (AFM)^39^ (Supplementary Fig. 3). Within the BTRR complex models, two predictions successfully identified the binding of the N-terminus of BLM to both TOP3A and the DUF1767 domain of RMI1 (Supplementary Figs. 3a; models 1 & 4; 3b; i), which aligns well with previously published experimental data^18,27,40,41^.

Two AF models indicated a direct BLM-PICH interaction, but unexpectedly, this was not via the PICH ABBA motif. Instead, this mapped between the core DNA-binding domain of PICH (SNF2-type helicase and HELICc domains) and one of the two RPA-binding (RB) motifs in BLM^5^ (Supplementary Figs. 3a; models 1 & 2; 3b; ii). Notably, all AF models confidently positioned the ABBA motif of PICH within the hydrophobic pocket of the OB-fold domains of the RMI1-RMI2 heterodimer^42,43^ (Fig. 2a; upper panel and Supplementary Figs. 3a models 1-5; 3b; iii). This RMI OB-fold dimer has been crystallised and shown to bind a different SLiM, MM2, found in the FANCM DNA translocase^23^ (Fig. 2a; bottom panel). Mutations in the MM2 motif compromise the recruitment of BTRR complexes to damaged replication forks^22^. Similar to the X-ray structure of MM2-RMI OB-fold dimers^23^, the ABBA motif of PICH was arranged under the same ‘knobs-in-holes’ organisation, in which alternating hydrophobic side chains of the motif project into hydrophobic pockets present on the composite surface made by both RMI1 and RMI2, albeit with opposite polarity (Fig. 2a). Highly conserved hydrophobic residues contributing to the interaction were noticed in both SLiMs. They include Phe953, Leu955 and Leu957 in PICH, and Phe1228, Phe1232, Leu1234 and Phe1236 in FANCM, respectively (Figs. 2a, 2b). In parallel, it is worth noting that AlphaFold models generated using full-length FANCM always predicted the opposite binding orientation of the MM2 motif as compared to the X-ray crystal structure^23^ (Supplementary Figs. 4a, 4b). Given the symmetrical arrangement of alternating hydrophobic residues in these SLiMs and their location in highly flexible regions, it is plausible that the interaction can occur bidirectionally. Thus, SLiM binding orientation may be influenced by the overall architecture of a protein complex in vivo.

**Figure 2.**
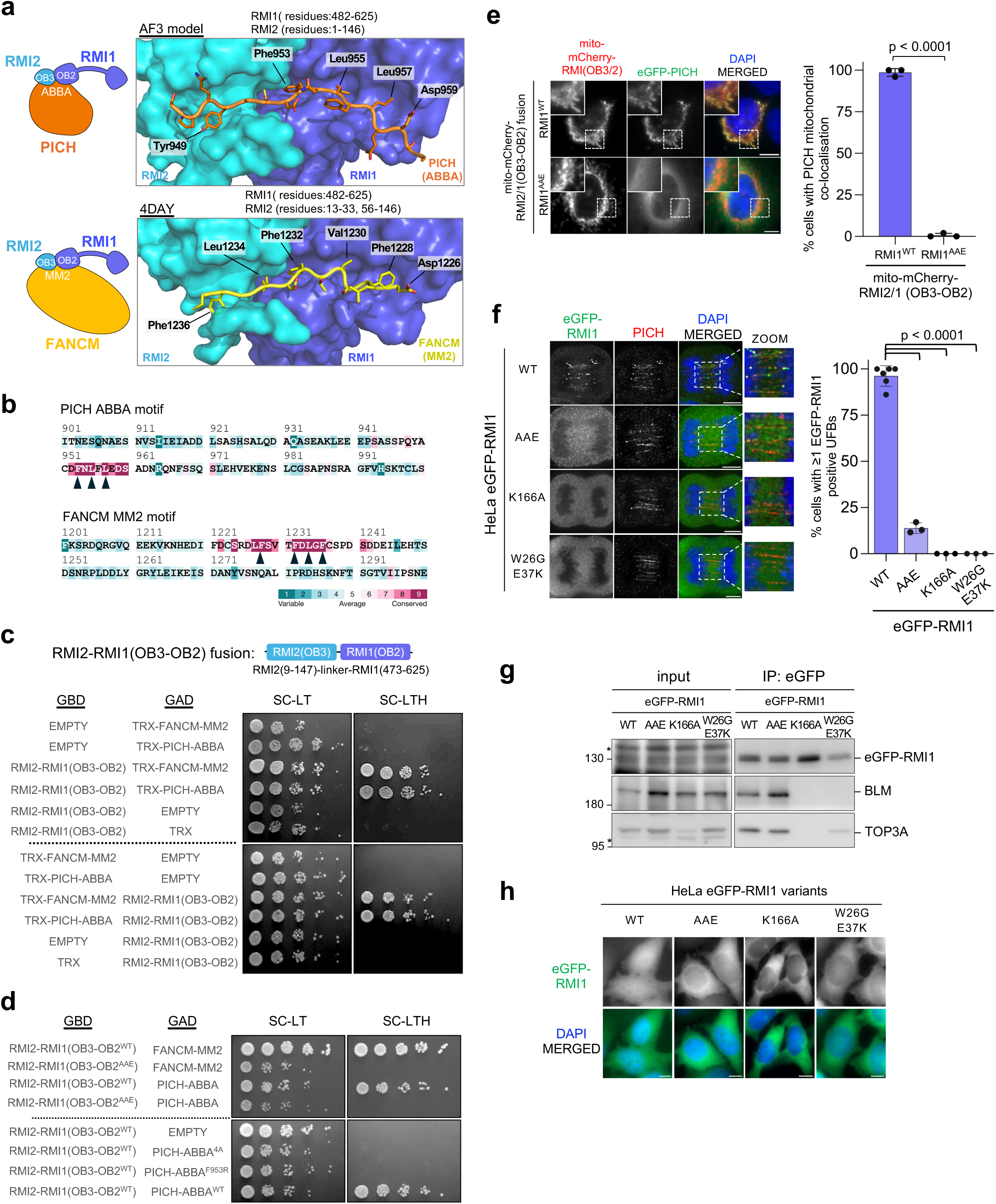
Both SLiM- and BLM-binding interfaces are essential for UFB association. **a.** Structural comparison of the RMI OB-fold pocket with the SLiMs of the PICH and FANCM DNA translocases. Upper: AlphaFold3 modelling of the PICH ABBA motif with the indicated regions of RMI1-RMI2. Bottom: X-ray crystal structure (PDB: 4DAY) of the FANCM MM2 motif with RMI1-RMI2. **b.** ConSurf protein sequence analysis of the SLiMs of PICH and FANCM. Arrowheads indicate the conserved hydrophobic residues contributing to interaction with the RMI OB-fold pocket. **c.** Yeast two-hybrid (Y2H) interaction analysis of the RMI OB-fold pocket with PICH- or FANCM-SLiMs (GBD: GAL4 DNA-Binding Domain; GAD: GAL4 Activation Domain). **d.** Mutations at the conserved residues of the RMI1 OB2 domain indicated in Supplementary Fig. 5**a** or at the PICH ABBA motif abolish their interaction, as shown by Y2H analysis. **e.** Mutations of the RMI1 OB2 domain abolish PICH interaction in cells, as determined by the MitoTrap PPI analysis. Left: Representative images of HeLa cells expressing mito-mCherry-RMI2-RMI1 OB-fold fusion and eGFP-PICH. Right: percentages of cells showing PICH mitochondrial localisation. Total number (n) of cells examined across three independent experiments: RMI1^WT^, n = 290; RMI1^AAE^, n = 301. **f.** UFB localisation of the indicated RMI1 variants on PICH-bound anaphase UFBs. Left: representative deconvolved images of the eGFP-tagged RMI1 variants. UFBs were revealed by PICH staining. Right: percentages of anaphase cells showing ≥ 1 eGFP-RMI1-bound UFB. Total number (n) of cells counted across at least three independent experiments: RMI1^WT^, n = 199; RMI1^AAE^, n = 109; RMI1^K166A^, n = 102; RMI1^W26G&E37K^, n = 95; mean ± S.D. is shown. Scale bars, 5 μm. **g.** Co-immunoprecipitation from asynchronous HeLa cells stably expressing the indicated eGFP-RMI1 variants to assess the effect of RMI1 mutations on BLM and TOP3A interaction. **h.** The effects of the indicated mutations on RMI1 nuclear localisation. Scale bars, 10 μm. The p-value is calculated by unpaired two-tailed t-test; Scale bars, 10 μm.

### The ABBA and MM2 motifs are ROB (RMI-OB-fold binding) motif

To validate the AlphaFold models of the UFB-binding complex, we first employed yeast two-hybrid (Y2H) assays. A fusion protein comprising the RMI2 OB3 and RMI1 OB2 domains was generated to reconstruct the putative SLiM-binding pocket and to test interactions with the PICH ABBA motif (Fig. 2c, top panel). The MM2 motif of FANCM was used as a positive control. Consistent with the AF predictions, the ABBA motif, like MM2, exhibited robust interaction with the RMI OB-fold pocket (Fig. 2c). Importantly, mutations of the conserved residues within the OB2 domain of RMI1 (Thr516Ala, Leu517Ala, and Leu578Glu; hereafter referred to as RMI1^AAE^ mutant) that are predicted to mediate SLiM interactions completely abolished binding to both ABBA and MM2 motifs (Fig. 2d and Supplementary Fig. 5a). Reciprocally, mutations of either the four conserved residues in the ABBA motif (FLFE to AAAA; referred to as 4A) or a substitution of phenylalanine 953 with arginine (F953R) also inactivated the interaction (Fig. 2d). Secondly, we examined the interaction between the RMI OB-fold pocket and full-length PICH proteins in human cells using MitoTrap. The OB-fold pocket fusion containing either RMI1^WT^ or RMI1^AAE^ was tagged with Mito-mCherry (Supplementary Fig. 5b). Consistent with the Y2H results, we observed that wildtype RMI1 but not the AAE mutant, was able to recruit full-length PICH to the mitochondria (Fig. 2e). These findings corroborate the structural models and show that the RMI OB-fold pocket can act as a molecular reader for SLiMs since it can recognise the hydrophobic-rich yet sequence-diverse motifs of two distinct SF2 DNA translocases, PICH and FANCM. We therefore designate the ABBA and MM2 motifs as the <u>R</u>MI <u>O</u>B-fold <u>b</u>inding (ROB) motif.

### The BTRR complex requires both the SLiM-binding pocket and BLM to engage with UFBs

To determine the importance of the RMI OB-fold pocket for the assembly of UFB-binding complexes on UFBs, we generated HeLa cell lines stably expressing eGFP-tagged RMI1 variants. Mutating the conserved residues in RMI1 (TLL to AAE) responsible for binding SLiMs significantly impaired RMI1 localisation to UFBs without affecting its BLM and TOP3A interactions (Figs. 2f, 2g). Similar to the incomplete loss of BLM association in PICH^ABBA^ cells, we observed that the RMI1^AAE^ pocket mutant exhibited residual UFB binding in a subset of anaphase cells. Given the AF predictions for potential BLM-PICH interactions, we wondered whether BLM and/or TOP3A may contribute to UFB association. We disrupted the interaction of RMI1 with both BLM and TOP3A through the RMI1 Lys166Ala (K166A) mutation^40,44^(Fig. 2g). Guided by AF modelling (Supplementary Fig. 5c), we also generated an RMI1 mutant, Trp26Gly and Glu37Lys (W26G/E37K), that loses the binding to BLM but not to TOP3A (Fig. 2g). Intriguingly, both the K166A and W26G/E37K mutants exhibited severe UFB localisation defects (Fig. 2f), indicating that BLM is essential for the stable loading of the TOP3A/RMI1/RMI2 (TRR) subcomplex onto DNA bridges, consistent with our recent work^27^. This finding contrasts with a previous report claiming that UFB association of the TRR subcomplex is BLM-independent^45^. On the other hand, we noticed that efficient nuclear import of RMI1 (and likely RMI2) also relies on BLM and/or TOP3A interactions because both K166A and W26G/E37K RMI1 mutants displayed a strong cytoplasmic localisation in interphase cells (Fig. 2h). Unexpectedly, we found that the RMI1 OB-fold pocket mutant, AAE, is also defective in nuclear import, albeit to a lesser extent than the BLM/TOP3A-binding mutant, K166A (Fig. 2h). This observation indicates that SLiM recognition by RMI1 has multifaceted functions, not only for interacting with different SF2 DNA translocases but also for facilitating efficient nuclear transportation of the RMI subcomplex.

### BLM secures BTRR complexes on DNA bridges through both PICH- and DNA-binding activities

Although recognition of the PICH ROB motif by RMI is crucial for stable engagement of the BTRR complex on UFBs, our data demonstrate that BLM is also indispensable. To investigate the molecular basis of this, we generated 25 AF-Multimer models of BLM and PICH. The predictions were classified into four groups based on interaction patterns. There are three potential sites in the N-terminal unstructured region of BLM exhibiting PICH-binding abilities (Supplementary Fig. 6). Notably, all models suggested that the second RPA-binding motif (RB-2) of BLM^5,46^ can interact with the core domains of PICH (Supplementary Fig. 6). We thus applied the MitoTrap PPI assay to validate the predictions. Since BLM is a nuclear protein, we removed its NLS (residues 1334-1417) to maintain the protein in the cytoplasm. To eliminate indirect binding to PICH through the RMI subcomplex, we also deleted the N-terminus of BLM (Fig. 3a). Consistent with AF predictions, only the amino- and not the carboxy-terminal half of BLM was able to drive mitochondrial recruitment of eGFP-PICH (Figs. 3a, 3b). Importantly, deletions of the region harbouring the RB-2 motif completely abolished PICH mitochondrial recruitment, supporting the proposed role of this motif for PICH interaction in cells. In agreement with this, we found that small BLM fragments containing the RB-2 motif were also sufficient to mediate PICH interaction (Figs. 3a, 3b). Reciprocally, we found that only the N-terminal part of PICH was able to interact with the N-terminal fragments of BLM (Figs. 3c, 3d). Therefore, the RB-2 motif can mediate binding both to RPA^5^ and to PICH. This finding may help explain the incompatibility between RPA and PICH binding by the BTRR complex observed in single-molecule experiments^45^.

**Figure 3.**
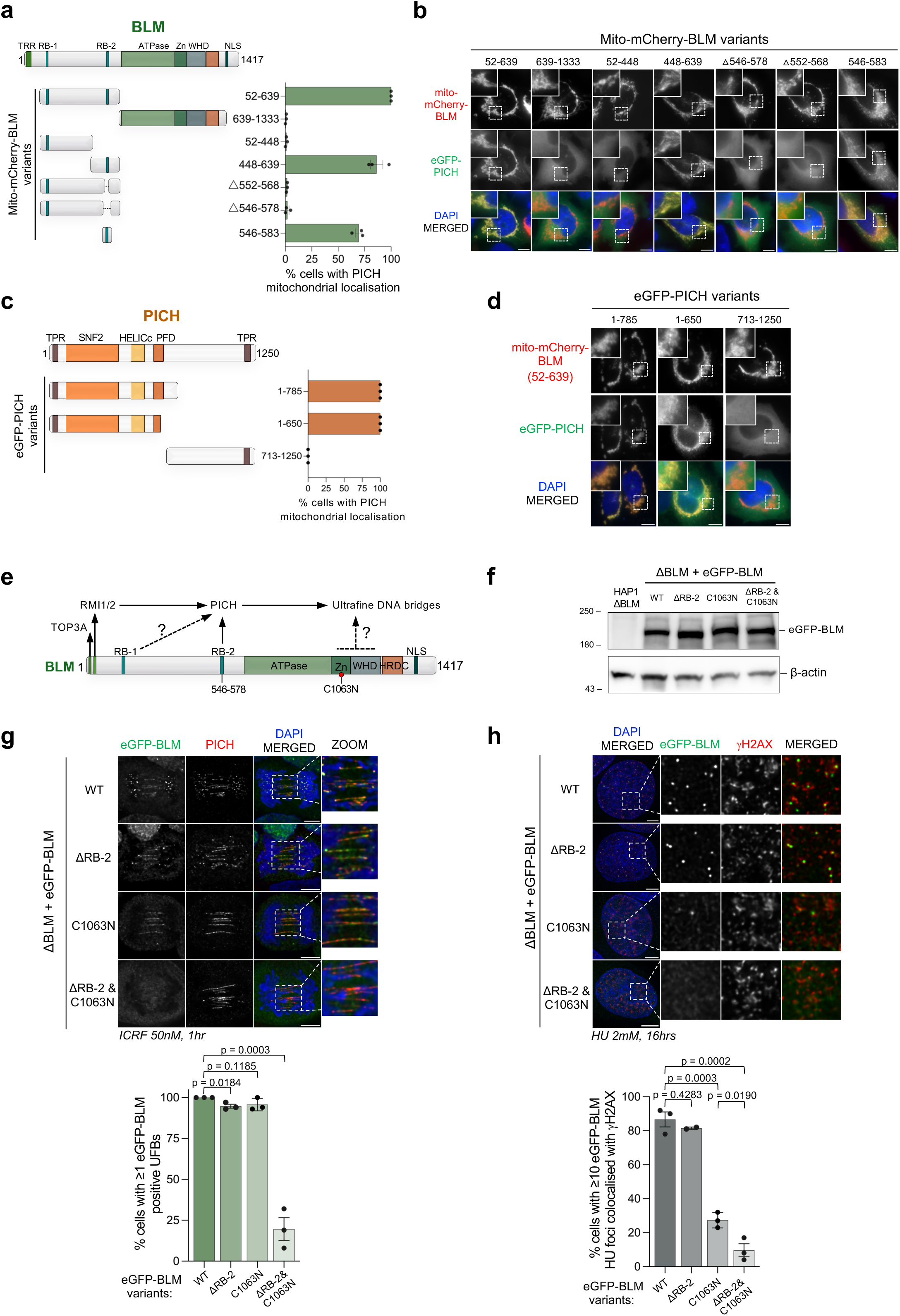
The PICH- and DNA-binding activities in BLM are essential for stable UFB engagement. **a.** MitoTrap PPI analysis of the indicated BLM truncated fragments for the interaction with full-length PICH. Domain organisation of BLM is shown (RB: RPA-binding motif; Zn: zinc finger domain; WHD: Winged-Helix domain; HRDC: Helicase-and-RNaseD C-terminal domain; NLS: Nuclear localisation signal). Right: quantification of cells showing positive PICH mitochondrial co-localisation. Total number (n) of cells analysed from three independent experiments: BLM fragments: 52-639, n = 303; 639-1333, n = 301; 52-448, n = 311; 448-639, n = 309; 52-639 Δ552-568, n = 303; 52-639 Δ546-578, n = 307; 546-583, n = 263; mean ± S.D. is shown. **b.** Representative images of MitoTrap PPI between the indicated BLM truncated fragments fused with Mito-mCherry and eGFP-tagged full-length PICH on mitochondria in HeLa cells. **c.** MitoTrap PPI analysis of the indicated PICH truncated fragments for the interaction with the BLM N-terminal fragment comprising residues 52-639. Right: Quantification of cells showing positive PICH mitochondrial co-localisation. Total number (n) of cells analysed from three independent experiments: 1-785, n = 503; 1-650, n = 368; 713-1250, n = 500; mean ± S.D. is shown. **d.** Representative images of MitoTrap PPI between the eGFP-tagged PICH truncations and Mito-mCherry-BLM (fragment 52-639) in HeLa cells. **e.** A diagram showing known (solid lines) and potential (dotted lines) sites of BLM for the interactions with the indicated proteins and anaphase bridge DNA molecules. **f.** Stable expression of the BLM variants tagged with eGFP in HAP1 ΔBLM cells. ΔRB-2: deletion of residues 546-578. β-actin acts as a loading control. **g.** UFB localisation of the stably expressed eGFP-tagged BLM variants on PICH-coated UFBs in HAP1 ΔBLM cells. Top: representative images of UFB staining. Bottom: Percentages of anaphase cells showing BLM-bound UFBs. Total number (n) of cells counted from at least three independent experiments: WT, n = 97; ΔRB-2, n = 99; C1063N, n = 96; ΔRB-2 & C1063N, n = 98; mean ± S.D. is shown. Scale bars, 5 μm. **h.** BLM localisation at hydroxyurea (HU)-induced DNA damage foci in cells expressing the indicated eGFP-BLM variants. Top: examples of deconvolved immunofluorescent images. Bottom: Percentages of cells showing ≥ 10 BLM foci colocalising with *γ*H2AX. Total number (n) of cells counted from at least three independent experiments: WT, n = 377 cells; ΔRB-2, n = 302 cells; C1063N, n = 451 cells; ΔRB-2 & C1063N, n = 398 cells; mean ± S.D. is shown. All p-values are calculated by unpaired two-tailed t-test. All scales bars, 10 μm (unless otherwise specified).

Next, we investigate whether the BLM-dependent loading of the BTRR complex to UFBs is driven by the RB-2 motif (Fig. 3e). We found that deleting the RB-2 motif did not compromise BLM UFB localisation (Figs. 3e-g). Possible explanations for this include a potential compensation mediated by the first RPA-binding (RB-1) motif. Alternatively, there might be other unidentified PICH-interacting site(s) for ancillary PPIs. We also wondered whether BLM’s DNA-binding activity is important for stabilising the BTRR complex-DNA interaction on UFBs (Fig. 3e). We favoured the latter possibility because a previous report showed that both RPA-binding motifs in BLM were dispensable for UFB localisation^5^. Additionally, our PPI analysis did not identify any other potential BLM-PICH interacting sites beyond the RB-2 motif. Thus, we decided to mutate the Zn-binding motif that has previously been shown to abolish BLM DNA-binding activity^47^. While the Zn-binding mutation, C1063N, alone did not impair BLM loading onto UFBs, combining it with deletion of the RB-2 motif resulted in a significant loss of UFB binding (Figs. 3f, 3g). Therefore, the BTRR complexes rely on either the BLM-PICH interaction or BLM-DNA binding to anchor stably on anaphase DNA bridges. These data explain why the interaction between BLM and the TRR subcomplex is critical. In contrast, we found that BLM binding to hydroxyurea (HU)-induced DNA lesions is highly dependent on the Zn-binding motif only (Fig. 3h), which indicates distinct determinants for the association with damaged replication forks and mitotic sister-DNA intertwining structures.

### PICH nuclear retention dampens DNA damage responses

That the ROB motifs of PICH and FANCM share the same binding pocket in the BTRR complex raises the possibility of molecular competition, but PICH is primarily cytoplasmic, whereas FANCM is nuclear. We speculated that nuclear export of PICH post-mitosis may be important to avoid its prolonged engagement with BTRR complexes, which would interfere with the BTRR complex functions during interphase. However, this hypothesis seemed inconsistent with a recent report proposing that a stable PICH-BTRR complex cannot form unless cells enter mitosis^48^. To address this, we examined the dynamics of PICH-BTRR complex formation in cells using the CenTrap PPI system (Fig. 4a). mCherry-tagged PICH was immobilised at centromeres through fusion to a truncated CENPB peptide^49^. We found that the CENPB-tagged PICH readily recruits different subunits of BTRR complexes to centromeres in interphase cells (Fig. 4b and Supplementary Fig. 7a) in a manner independent of PICH activity but partially dependent on the ROB motif (Supplementary Figs. 7b, 7c). This indicates that the PICH-BTRR complex assembly remains competent during interphase, regardless of whether it is in the nucleus or cytoplasm. Interestingly, we observed that as cells enter mitosis, this complex formation rapidly diminishes but is restored upon the onset of anaphase (Fig. 4b and Supplementary Fig. 7d). These results align well with our previous proposal of phosphorylation-dependent inhibitory regulation in early mitosis^27^. Notably, a similar phenomenon was detected between FANCM and the BTRR complex, albeit with less obvious reassembly after anaphase onset (Fig. 4b). We also examined nuclear interaction between PICH and RIF1. In contrast to the BTRR complex, centromeric recruitment of RIF1 by the CENPB-tagged PICH was less efficient and only observed in a subset of cells, suggesting that their interaction may be weaker. Nevertheless, this is still mediated by PICH ROB motif (Supplementary Fig. 7e).

**Figure 4.**
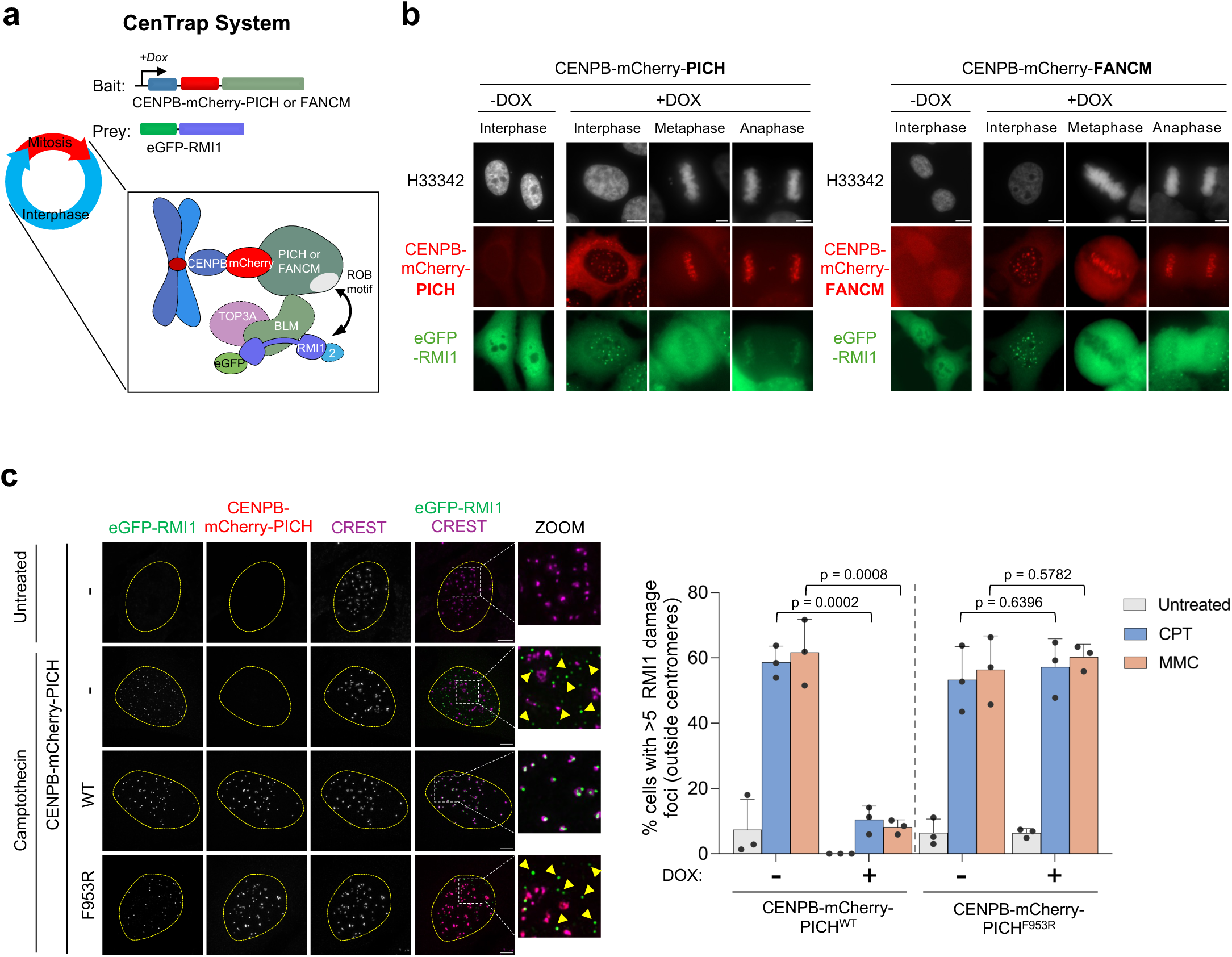
Nuclear immobilisation of PICH inactivates BTRR complexes in response to replication fork damage. **a.** The CenTrap Protein-Protein Interaction (PPI) System to test interactions between eGFP-tagged RMI1 and PICH or FANCM through the cell cycle. **b.** Representative images of the centromeric localisation of CENPB-mCherry PICH/FANCM and eGFP-tagged RMI1 in the indicated phases of the cell cycle. **c.** Immunofluorescence microscopy showing HeLa cells with or without the expression of CENPB-mCherry-PICH^WT^ or PICH^ABBA^ following the indicated treatments. Centromeres are labelled by CREST. Arrowheads indicate DNA damage foci marked by eGFP-RMI1 localised outside the centromeric regions. Right: quantifications of the indicated cells displaying eGFP-RMI1 DNA damage foci after camptothecin (CPT) or mitomycin C (MMC) treatments. Total number (n) of cells counted from three independent experiments: PICH^WT^ Dox (-) UT, n = 353; CPT, n = 353; MMC, n = 358; PICH^WT^ Dox (+) UT, n = 306; CPT, n = 340; MMC, n = 318; PICH^F953R^, Dox (-) UT, n = 401; CPT, n = 375; MMC, n = 362; PICH^F953R^, Dox (+) UT, n = 305; CPT, n = 316; MMC, n = 307; mean ± S.D. is shown. All p-values are calculated by unpaired two-tailed t-test; Scale bars, 10 μm.

Next, we tested the impact of inappropriate engagement of the PICH-BTRR complex in the interphase nucleus on the functions of BTRR complexes, particularly on DNA damage responses. We treated cells with camptothecin (CPT) or mitomycin C (MMC), as both induce BTRR damage foci in a FANCM-dependent manner^22^. As expected, CPT and MMC treatments readily induced DNA damage foci labelled by ectopically expressed eGFP-RMI1, which predominantly appeared outside centromeric regions (Fig. 4c). Expressing the nuclear-targeted PICH fusion (CENPB-mCherry-PICH) largely suppressed the formation of the eGFP-RMI1 damage foci. This suppression was dependent on the PICH ROB motif (Fig. 4c and Supplementary Fig. 7f). Thus, persistent occupancy of the RMI OB-fold pocket by PICH can attenuate the response of BTRR complexes to DNA damage in interphase cells. Therefore, we propose that, in addition to mitotic phosphorylation inhibition^26,27^, active nuclear exclusion of PICH may serve as another important regulatory mechanism to prevent illegitimate PICH-BTRR complex engagement.

### Multiple nuclear export activities counteract PICH nuclear retention after mitosis

Although PICH shares significant sequence homology with the common ancestors of chromatin remodelers, including BTAF1, ERCC6 and RAD54 (Supplementary Fig. 8), it lacks a canonical nuclear localisation signal (NLS)^50,51^ and is actively excluded from the nucleus until the onset of mitosis. Typically, nuclear export of proteins is driven by the Exportin 1 (XPO1)-dependent pathway^52^. Consistent with this, AF predictions strongly indicate that a short helix within PICH (residues 765-777) directly engages XPO1 (Fig. 5a). Although the amino acid sequence of this helix shows moderate conservation across species, it closely resembles the consensus leucine-rich nuclear export signal (NES) characterised in SNUPN^52^ (Figs. 5a, 5b). Moreover, treatment with the XPO1 inhibitor, leptomycin B (LMB), compromised the cytoplasmic localisation of PICH. However, this effect was predominantly restricted to newborn G1 cells (Fig. 5c; arrows), indicating that its nuclear localisation was likely due to PICH being trapped in the reforming nucleus rather than to active nuclear import. A systematic series of PICH truncations validated the AF predicted helix as a bona fide NES (Fig. 5d; i-viii). Unexpectedly, deletion of this primary NES only partially weakened the nuclear exclusion of PICH (Fig. 5d; ix), strongly indicating the presence of additional, redundant NES(s). Supporting this, we identified two other regions within PICH that exhibit autonomous nuclear export activity: one mapped within the core domains, and another located at the C-terminus between residues 900 and 1000 (Fig. 5d; xiii-xviii). The latter region coincides with the ROB motif, and mutations within this sequence partially compromised nuclear exclusion (Fig. 5d; xvi-xviii). Interestingly, although in silico analyses failed to predict a canonical NLS in PICH, we identified two distinct regions displaying intrinsic nuclear localisation activity (Fig. 5d; x-xii; blue boxes). Nevertheless, these nuclear import activities appear to be overridden by the multiple, robust NES modules identified here. Taken together, PICH may have undergone evolutionary optimisation to degenerate classical NLSs and acquire strong nuclear export activity to execute yet-unidentified cytoplasmic functions and/or to prevent unnecessary interaction with the BTRR complex that could disturb the DNA damage response.

**Figure 5.**
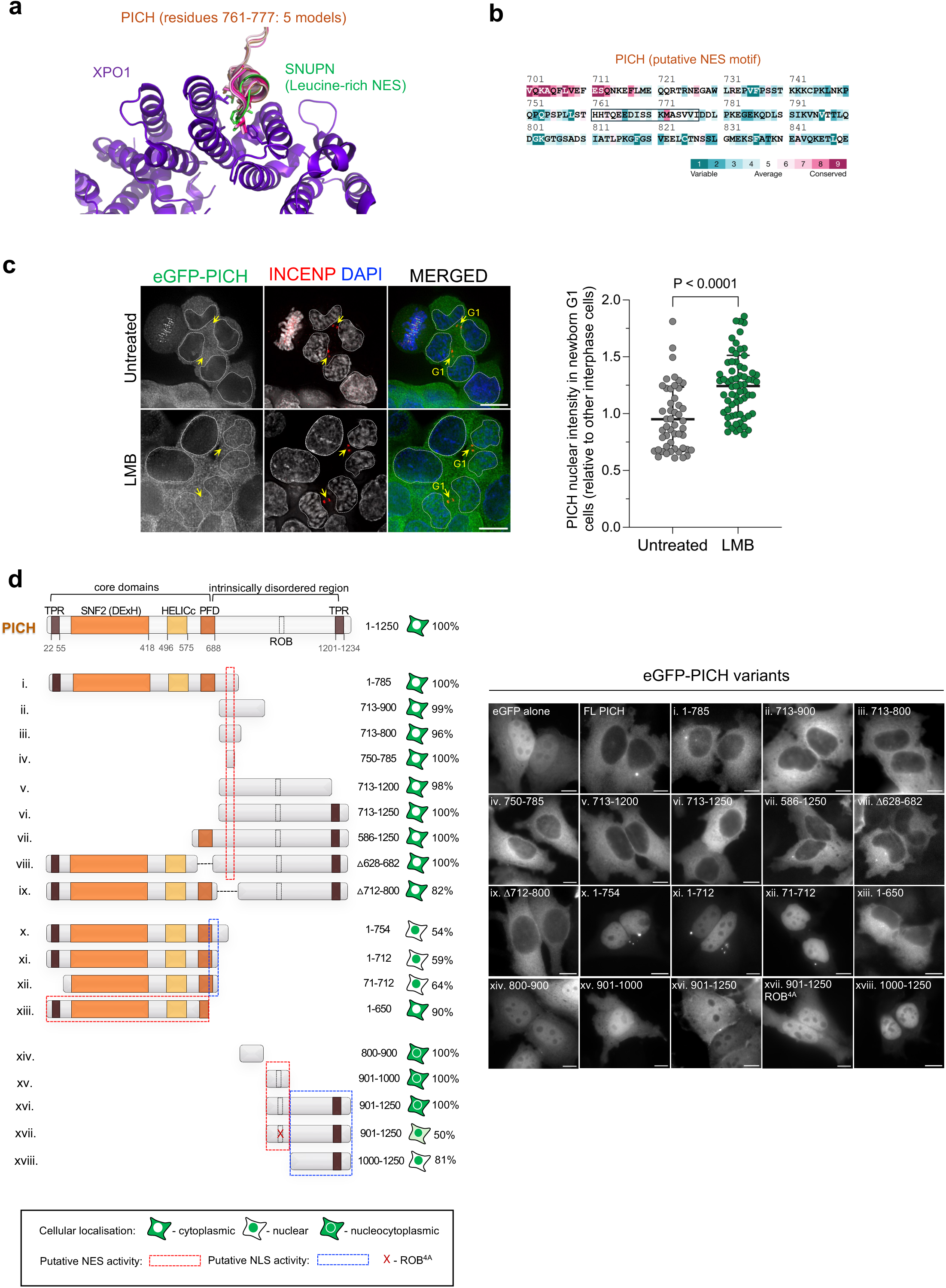
PICH possesses three modules that exhibit nuclear export activity. **a.** AlphaFold3 prediction of the interaction between PICH and XPO1. Five models indicate that PICH residues 761-777 bind to the same site in XPO1 bound by SNUPN leucine-rich NES (PDB: 3GB8). **b.** ConSurf analysis shows moderate conservation of PICH residues 761-777. **c.** Short-term inactivation of XPO1 results in PICH nuclear retention in newborn G1 cells. Left: immunofluorescence microscopy showing HCT116 cells stably expressing eGFP-PICH treated with and without the XPO1 inhibitor, leptomycin B (LMB), for 2 hours. Newborn G1 cell pairs are identified by INCENP staining (yellow arrowheads). Right: measurements of PICH nuclear intensity in newborn G1 cells compared with that in other interphase cells. Total number (n) of cells analysed: untreated: INCENP-negative cells, n = 72; INCENP-positive newborn G1 cells, n = 49; LMB-treated: INCENP-negative cells, n = 100; INCENP-positive newborn G1, n = 66; mean ± S.D. is shown. **d.** Identification of domains in PICH exhibiting nuclear export and import activities. Systematic truncation analysis of the indicated PICH variants showing nuclear or cytoplasmic localisations. Left: red dotted regions indicate nuclear export activity, whereas blue dotted regions show nuclear localisation activity. Right: representative fluorescent images of the cellular localisation of the eGFP-PICH variants in HeLa cells. eGFP alone is used as a reference. Scale bars, 10 μm.

## Discussion

In this study, we present a comprehensive analysis of the molecular organisation of the UFB-binding complex. This leads us to uncover a conserved mechanism that enables BTRR complexes to engage distinct SF2 DNA translocases during DNA repair and chromosome segregation. This mechanism relies on the hydrophobic pocket formed in between the OB-fold domains of the RMI1-RMI2 heterodimer. Previously, it was shown that the RMI OB-fold domains can bind to the MM2 motif of the FANCM SF2 DNA translocase^22,23^, which is crucial for the recruitment of the BTRR complex to damaged replication forks^22^. Here, we demonstrate that PICH, another evolutionarily distant SF2 DNA translocase^24^, exploits the same mechanism to secure the BTRR complex loading onto UFB structures in human anaphase cells. Assembly of the BTRR complex with either FANCM or PICH is highly dependent on the ROB motifs embedded within the intrinsically disordered regions of these proteins. Despite their limited sequence identity, both ROB motifs contain multiple hydrophobic residues arranged in a similar pattern. We therefore propose that the RMI1-RMI2 heterodimer functions as a molecular reader of hydrophobic-rich SLiMs. Interestingly, the PICH ROB motif is also essential for RIF1 loading on DNA bridges. Whether RIF1, alone or in complex with PP1^48^, can form a structure similar to the RMI OB-fold pocket remains undetermined.

Although the SLiM-recognition system is indispensable, stable anchoring of the BTRR complex to UFBs still requires BLM. This finding is consistent with the general properties of SLiMs, which typically mediate weak and transient protein-protein interactions (PPIs) owing to their short length, often comprising fewer than ten amino acids^53^. Such dynamic interactions may nevertheless provide a mechanism for fine-tuning protein assemblies, particularly when combined with additional low-affinity interfaces. Indeed, we find that this additional binding support is provided by the RPA-binding motif 2 (RB-2) or DNA-binding activity of BLM. The latter finding is perhaps surprising, as the prevailing view is that PICH acts as the primary UFB sensor, assembling the entire UFB-binding complex primarily through protein-protein interactions. Collectively, our findings reveal a mechanistic model for UFB-binding complex formation on intertwined DNA bridge structures (Fig. 6). Upon the onset of anaphase, dimeric or multimeric PICH regains the ability to interact with both the BTRR and RIF1-PP1 complexes^25,27^. The BTRR complex exploits BLM-PICH interactions (Fig. 6; 1), BLM-mediated DNA binding (Fig. 6; 2), and the RMI-dependent SLiM recognition mechanism (Fig. 6; 3) to anchor stably to the PICH-bound DNA bridges, promoting DNA unwinding. This process may require initial chromatin remodelling by PICH to generate a DNA substrate for BLM. Indeed, a recent report has shown that the ATPase activity of PICH is needed for BLM-mediated UFB unwinding, albeit not for BLM recruitment^37^. In contrast, recruitment of RIF1-PP1 may be driven primarily by binding to the PICH ROB motif (Fig. 6; 4) because mutations of PICH ROB motif completely abolish RIF1’s UFB localisation.

**Figure 6.**
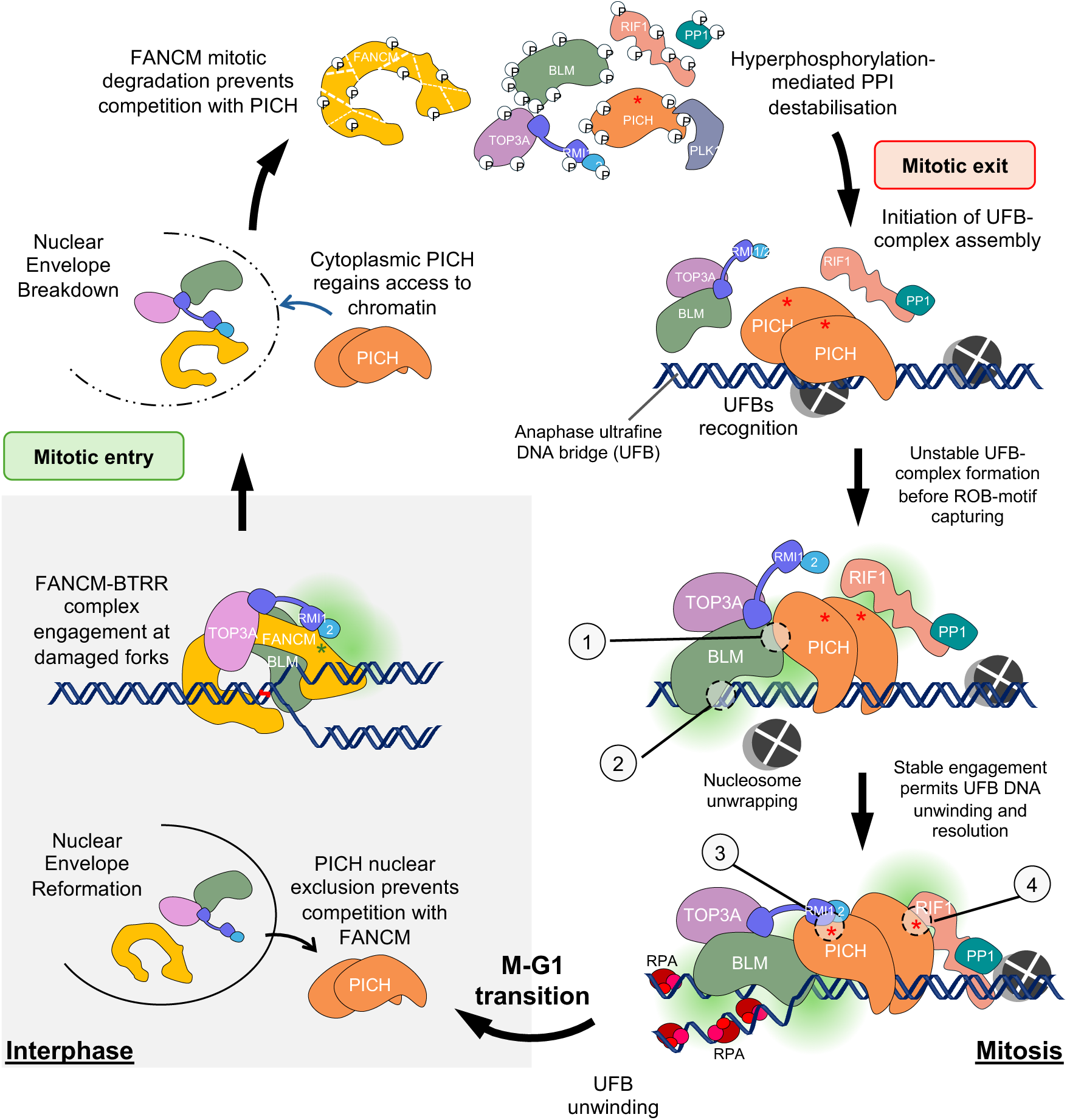
A model of the dynamic assembly of the UFB-binding complex for genome stability maintenance. Following mitotic entry and nuclear envelope breakdown, cytoplasmic PICH gains access to chromatin. Mitotic hyperphosphorylation transiently restricts stable assembly of PICH, the BTRR complex and RIF1-PP1 to protect centromere chromatin. FANCM undergoes mitotic protein degradation, minimising subsequent competition with PICH for the BTRR in late mitosis. At the onset of anaphase (mitotic exit), the inhibitory phosphorylation is removed, enabling both the BTRR complex and RIF1-PP1 to associate with PICH along the ultrafine DNA bridges (UFBs). The BTRR complex exploits RB-2-mediated PICH interactions (1), BLM-mediated DNA binding (2), and the RMI-dependent SLiM recognition (3; red asterisk) to stably anchor onto the PICH-bound DNA bridges, promoting DNA unwinding and subsequent RPA loading. The latter might require PICH-mediated nucleosome remodelling. In contrast, recruitment of RIF1-PP1 is presumably mediated only through the interaction of the PICH ROB motif (4; red asterisk). Following UFB resolution, the UFB-binding complexes disassemble, and the PICH DNA translocase is rapidly excluded from the nucleus after nuclear envelope reformation during the M-to-G1 transition, thereby preventing persistent occupancy of the RMI OB-fold pocket. This permits the BTRR complex to interact with other ROB motif-containing proteins, such as FANCM (green asterisk), and act during the repair of damaged replication forks until the entry into the next mitosis.

Given its stoichiometric organisation, each BTRR complex is expected to bind only a single DNA translocase at any given time, assuming no additional DNA translocase-binding sites are available. However, as shown here and by others, both PICH and BLM can form dimers or high-order oligomers^46,54,55^, which raises the possibility that multiple DNA translocases could associate with BTRR complex multimers and vice versa. Although speculative, such higher-order configurations may be required during complex chromatin remodelling and the restart of stalled replication forks. Nevertheless, it is highly plausible that the UFB-binding complex exists as a large supramolecular protein structure on anaphase bridges. Conversely, timely disassembly of the UFB-binding complex is equally important. As shown here, persistent occupancy of the RMI OB-fold pocket by PICH can interfere the DNA damage response of the BTRR complex. We previously demonstrated that premature assembly of the UFB-binding complex is suppressed by CDK1 and PLK1 in early mitosis for centromere protection^26,27^. However, an equivalent inhibitory control appears to be absent following mitotic exit. To prevent illegitimate engagement of the PICH-BTRR complex post-mitosis, we propose that active nuclear export of PICH provides an additional regulatory layer. Consistent with this notion, the nuclear localisation signals (NLSs) within PICH appear to have substantially degenerate during evolution^50^, whereas multiple nuclear export modules have evolved. Although a previous study proposed an interphase role of PICH for recovery of stalled replication forks^56^, it remains unclear how cells would simultaneously overcome robust nuclear export and activate nuclear translocation of PICH. Alternatively, a very small pool of PICH may be retained in interphase nucleus. Nevertheless, our findings suggest that efficient exclusion of PICH following nuclear envelope reformation minimises unnecessary sequestration of the BTRR complex, thereby preserving DNA damage response capacity. This model provides a possible explanation for the longstanding observation that PICH predominantly localises to the cytoplasm^24^, although additional cellular functions cannot be excluded.

Reassembly of cytoplasmic PICH with nuclear BTRR complexes occurs following nuclear envelope breakdown, particularly after the removal of mitotic phosphorylation in anaphase^26,27^. In principle, FANCM could compete with PICH for the BTRR complex during mitosis, perturbing the loading onto the UFBs. This could be potentially prevented via mitotic phosphorylation, as we observe FANCM-BTRR complex dissociation during mitosis. More interestingly, a previous study found that FANCM undergoes rapid protein degradation upon mitotic entry through a PLK1- and β-TRCP-dependent pathway^57^, which could eliminate competition between FANCM and PICH during late mitosis (Fig. 6).

On the other hand, the formation of the FANCM-BTRR complex plays a critical role in telomere maintenance in alternative lengthening of telomeres (ALT) tumour cells^58^. Efforts have been devoted to developing small-molecule inhibitors and peptide mimetics that target the RMI SLiM-binding pocket as potential anti-cancer therapeutics^59,60^. However, our findings raise an important concern about whether such inhibitors may also disrupt interactions with other protein(s) that possesses ROB motifs, including PICH and factors that remain unidentified. Unexpectedly, we have also uncovered a role for the RMI OB-fold pocket in the nuclear import of RMI proteins. This observation suggests that SLiM recognition by RMI may extend functions beyond DNA repair and chromosome segregation. Therefore, the potential effects of therapeutic strategies targeting the SLiM-binding pocket should be further explored.

In summary, we define the molecular architecture of the UFB-binding complex assembled on sister DNA intertwinements in human anaphase cells and uncover the mechanism that enables the spatiotemporal association among three distinct SF2 DNA helicases: BLM, as part of the BTRR complex, PICH, and FANCM. The dynamic exchange of PICH and FANCM with the BTRR complex is highly coordinated through cellular compartmentalisation, post-translational phosphorylation, and programmed protein degradation.

## Methods

### Plasmids and cloning

To mutate the ABBA-like motif of PICH at the endogenous locus, sgRNAs targeting PICH were cloned into a modified version of the PX466 ‘All-in-One’ plasmid^61^ containing Cas9D10A-T2A-mRuby via restriction digestion. The donor template (pAAV-PICH(ABBA)-repair-template) containing PICH ABBA mutations and flanking homology arms was custom-synthesised (Integrated DNA Technologies). Plasmids pMA-RQ-PICHmAID-GFP-Zeo and pX459-PICH-sgRNA for endogenous tagging of PICH with -mClover3 were gifts from Gary Ying Wai Chan. The mAID sequence was removed by restriction digestion. Plasmids expressing GFP-tagged RMI1 were constructed by subcloning human RMI1 cDNA (a gift from Weidong Wang) into the pSBbi-Neo vector (a gift from Eric Kowarz; Addgene plasmid #60525). Plasmids expressing eGFP-RMI1 and eGFP-BLM were subsequently mutagenised using the QuikChange XL Site-Directed Mutagenesis Kit (Agilent Technologies; #200516) or modified using the Gibson Assembly Master Mix (New England Biolabs; #E2611L) according to the manufacturers’ protocols.

To generate constructs for the MitoTrap assays, mito-mCherry was cloned into pSBtet-BN (Addgene plasmid # 60502) from pMito-mCherry-FRB_IRES_FKBP(III)-GBPen (a gift from Stephen Royle; Addgene plasmid # 128268) and different fragments of BLM and PICH were inserted via restriction digestion.

For the CenTrap assay, the mCherry-tagged DNA-binding domain of CENPB (a gift from Mark Burkard) was cloned into pSBtet-BN (Addgene plasmid #60496). Full-length human PICH and FANCM (a gift from Daniel Durocher) cDNAs were similarly inserted by restriction digestion. Truncated variants of PICH were generated by PCR amplification and directionally cloned into the pEGFP-C1 vector using restriction digestion. All oligonucleotides and primers used for cloning, mutagenesis, and sequencing are detailed in Supplementary Data 4.

### Cell culture and cell lines

hTERT RPE1 and RPE1 FRT/TR cells were cultured in DMEM/F-12 medium (Sigma) supplemented with 10% FCS (Gibco) and 1% Penicillin/Streptomycin antibiotics. HeLa and HCT116 cell lines and their derivatives were cultured in DMEM (Sigma) supplemented with 10% foetal calf serum (FCS) (Gibco), 1% L-glutamine, and 1% Penicillin/Streptomycin antibiotics. HAP1 and its derivatives are cultured in IMDM (Gibco) medium (Sigma) containing 10% FCS (Gibco) and 1%Penicillin/Streptomycin antibiotics. All human parental cell lines were obtained from the Cell Bank at the Genome Damage and Stability Centre, which verified them using ATCC’s cell line authentication service. All cell lines were periodically tested and confirmed to be free of mycoplasma (Lonza Mycoplasma testing kit). All cell lines were meticulously maintained at 37°C in a humidified 5% CO_2_ atmosphere. RPE1 FRT/TR PICH^ABBA^ mutant cell lines were generated by transfecting ‘All-in-One’ plasmid containing Cas9^D10A^-T2A-mRuby^61^ and gRNAs targeting around ABBA-like motif (5’-ATGTGATTTCAATCTTTTCT-3’ and 5’-TACTGTGGTGAAGATGCTGA-3’) along with the donor plasmid pAAV-PICH^ABBA^ repair template with F953A, L955A, F956A, and E958A using 1:5 ratio into cells. Transfection was performed by electroporation using the Neon Transfection Kit (ThermoFisher; MPK1025) at 1400 V for 20 ms with 2 pulses. Two days post-transfection, cells were sorted (BD FACSAria^TM^) based on mRuby expression and expanded into single colonies. Edited clones were identified by PCR using the forward primer 5’-CAAAGGGGTTTGGAAGTGTAGA-3’ and the reverse primers 5’-ACATGCATACTGTGGTGAAGA-3’ or 5’-GGCGTACTGGGGACTGGA-3’ (for wild type or mutant, respectively) and confirmed by sequencing (GeneWiz).

HeLa, HCT116 and HAP1 stable cell lines were generated by lipid-mediated transfection with plasmids encoding the gene of interest using FuGENE HD (Promega; TM328) according to the manufacturer’s guidelines. A BD FACS Melody^TM^ cell sorter was used to sort and generate stable cell lines expressing the indicated eGFP- or mCherry-tagged fluorescent proteins.

For ultrafine DNA bridge (UFB) localisation and centromere unwinding analyses, cells were first synchronised via a single thymidine block for 16 hours. They were then released into fresh, thymidine-free media for 6-8 hours to enrich G2/M populations. The indicated small-molecule inhibitors or drugs were added to the culture medium during the final 2 hours before cell fixation. For the MitoTrap assay, HeLa cells were seeded in 6-well plates and, after 24 hours, co-transfected with the indicated expression plasmids using FuGene. At 24 hours post-transfection, cells were harvested, reseeded onto glass coverslips, and incubated for a further 24 hours to allow recovery. To induce expression of pSBtet-mito-mCherry, cells were treated with doxycycline for 4-6 hours. Cells were then fixed with 3.7% paraformaldehyde (PFA), stained with Hoechst (H33342) for nuclear staining, and imaged.

For the CenTrap assay, HeLa cells stably expressing eGFP-RMI1 and inducible CENPB-mCherry-PICH were seeded onto glass coverslips and allowed to adhere for 6–8 hours. To induce expression of CENPB-mCherry-PICH, cells were treated with doxycycline. After 2–4 hours of induction, the DNA-damaging agents camptothecin (10 nM) and mitomycin C (40 ng/mL) were added to the culture medium, and cells were incubated for 16 hours. Cells were then fixed with 4% paraformaldehyde (PFA) and processed for immunofluorescence (IF).

### Chemicals and small molecule inhibitors

Doxycycline (Merck; D5207, 0.25-1 µg/ml) was used to induce expression of the gene of interest via the Sleeping Beauty (SB) transposon system. Thymidine (Sigma; T9250, 2 mM) and nocodazole (Sigma; SML1665; 50 ng/ml) were used to synchronise the cell cycle and arrest cells. ICRF193 (Merck; I4659; 30-50 nM) was used to induce UFBs. BI2536 (Cayman Chemical; 17385; 60-100 nM) was used to inhibit PLK1 kinase. Rapamycin (Sigma; R878; 200 nM) was used to dimerise Mito-mCherry-FRB and FKBP12-GFP binding protein in the MitoTrap assay testing PICH dimerisation. Mitomycin C (Roche; 1010740900140) and camptothecin (Sigma; 208925) were used to induce DNA damage. Leptomycin B (Sigma; L2913; 25nM) was used to inhibit XPO1 and block the nuclear export of PICH.

### Fluorescence-activated cell sorting (FACS)

Cells were harvested by trypsinisation, washed with 1× PBS, and fixed in 70% ice-cold ethanol. Fixed cells were resuspended in a propidium iodide (PI)/RNase A staining solution containing 40 μg/mL PI and 100 μg/mL RNase A in 1× PBS. Cell cycle profiles were acquired using a CytoFLEX flow cytometer (Beckman Coulter) and analysed using CytExpert software.

### Immunofluorescence staining

For immunostaining analysis, cells were seeded onto No. 1.5 or 1.5H cover glass and treated as indicated. Cells were fixed with Triton X-100-PFA buffer (250 mM HEPES, pH 7.4, 1× PBS, 0.1% Triton X-100, 4% methanol-free paraformaldehyde) at 4 °C for 20 min, or with PBS–PFA buffer (1× PBS, 3.7% methanol-free paraformaldehyde) at room temperature for 10 min. Fixed cells were washed with PBS, permeabilised with 0.1% Triton X-100 and 0.5% FCS in PBS for 20 min at 4 °C, then blocked (1× PBS, 0.5% FCS) at room temperature for 10 min. Cells were incubated with the primary antibody at 37 °C for 90 min, followed by incubation with the secondary antibody at room temperature for 30 min. Slides were then washed with 1× PBS and ultra-pure water, 3-4 times and once, respectively. Coverslips were air-dried before mounting with Vectashield mounting medium, either with or without DAPI. DAPI was omitted when DNA was stained with Hoechst H33342 (Invitrogen; C10637-G) prior to mounting. Fluorescence images were acquired on a Zeiss AxioObserver Z1 epifluorescence microscope equipped with 40x/1.3 oil Plan-Apochromat, 63x/1.4 oil Plan-Apochromat and 100x/1.4 oil Plan-Apochromat objectives and a Hamamatsu ORCA-Flash4.0 LT Plus camera. Z-stack images were acquired at 0.2 μm intervals over a depth range of 2–8 μm using ZEN blue software. Image deconvolution was performed using Huygens Professional with a measured point-spread function (PSF) obtained from 200-nm-diameter TetraSpeck microspheres (Thermo Fisher). The classical maximum likelihood estimation method was applied with 40-60 iterations and a signal-to-noise ratio of 20-60. Fiji was used to generate the representative images.

Primary antibody dilutions: goat anti-BLM (Santa Cruz; sc-7790), mouse anti-BLM (Santa Cruz; sc-365753), rabbit anti-BLM (Abcam; ab2179, 1:100), human anti-CREST (Immuno Vision; HCT-0100, 1:400), mouse anti-phospho-Histone H2A.X (Ser139) (Sigma Aldrich; 05-636, 1:400), mouse anti-PICH (Abnova; H00054821-BO1P, 1:100), rabbit anti-PICH (Abnova; H00054821-DO1P, 1:100), rabbit anti-RIF1 (Bethyl Lab; A300-568A, 1:100), rabbit anti-RIF1 (Bethyl Lab; A300-569A, 1:100), rabbit anti-RPA70 (Abcam; ab79398, 1:200); Secondary antibody dilutions: donkey anti-goat AF488 (Invitrogen; A-11055, 1:500), donkey anti-mouse AF488 (Invitrogen; A-21202, 1:500), donkey anti-rabbit AF488 (Invitrogen; A-21206, 1:500), donkey anti-mouse AF555 (Invitrogen; A-31570, 1:500), donkey anti-rabbit AF555 (Invitrogen; A-31572, 1:500), donkey anti-mouse AF647 (Invitrogen; A-31571, 1:500), donkey anti-rabbit AF647 (Invitrogen; A-31573, 1:500) and goat anti-human AF650 (Abcam; ab98622, 1:500). UFB intensity measurements: UFBs were stained for UFB-binding complex components, including BLM and PICH, following the immunofluorescence protocol. Fiji was used to measure the intensities of individual UFBs by drawing a scanline along the entire length of each UFB thread, using PICH staining as the reference. Channel intensities were corrected for background before further analysis. Final values were normalised to the average intensities measured in PICH^WT^ cells.

For nuclear intensity measurements of PICH, cells were categorised into newborn G1 phase (INCENP-positive) and non-G1 interphase (INCENP-negative) populations. The nuclear boundary of each cell was manually outlined based on the DAPI channel using the freehand selection tool in Fiji. The mean fluorescence intensity of eGFP-PICH within the defined nuclear area was then determined. Final values were normalised to the average PICH intensity measured in non-G1 interphase control cells.

### Western Blot

Cells were trypsinised and washed once with ice-cold PBS, then lysed in Lysis Buffer (50 mM Tris-HCl pH 7.5, 150 mM NaCl, 1% Triton X-100, 1.25 mM DTT, 1 mM PMSF, 25 U/ml Benzonase (Sigma-Aldrich; E1014)) on ice. The lysis buffer was supplemented with the Protease inhibitor cocktail cOmplete (Roche; 11697498001), 1 tablet/50 ml, and, when required, the phosphatase inhibitor PhosSTOP (Roche; 4906845001), 1 tablet/10 ml. Cell lysate was incubated on ice for 15-20 min and centrifuged at 15000 rpm for 20 min at 4°C. Protein concentrations were quantified using a Bradford assay (Bio-Rad). Samples were separated by SDS-PAGE, transferred onto Amersham Hybond PVDF membranes (0.2 *μ*m, GE Healthcare Life Sciences, 10600021), and blotted with the indicated antibodies following standard procedures.

Primary antibodies: mouse anti-β-actin (Sigma; A5441, 1:5000), goat anti-BLM (Bethyl Lab; A300-120, 1:2000), rabbit anti-BLM (Abcam; ab2179, 1:500), rabbit anti-GFP (Abcam; ab290, 1:2000), rat anti-GFP (Chromotek; 3h9-100, 1:2000), rabbit anti-Ku80 (Abcam; ab80592, 1:5000), rabbit anti-mCherry (Abcam; ab183628, 1:1000), mouse anti-PICH (Abnova; H00054821-BO1P, 1:500), mouse anti-PLK1 (Sigma-Aldrich; 05-844, 1:1000), rabbit anti-RIF1 (Bethyl Lab; A300-568A, 1:1000), rabbit anti-RIF1 (Bethyl Lab; A300-569A, 1:1000), rabbit anti-TOP3A (Proteintech; 14525, 1:500). HRP-conjugated secondary antibodies: rabbit anti-goat (Agilent; P0160, 1:60000), goat anti-mouse (Abcam; ab6780, 1:25000), donkey anti-rabbit (ECL/Sigma; NA9340, 1:40000), and goat anti-rat (ECL/Sigma; GENA935, 1:25000).

### Co-immunoprecipitation

Endogenous mClover3-tagged PICH proteins were immunoprecipitated from wild-type and ABBA-mutant PICH-mClover3 RPE1 FRT/TR cells. Cells were lysed in ice-cold lysis buffer (50 mM Tris-HCl, pH 7.5, 150 mM NaCl, 1% Triton X-100, 25 mM DTT, 1 mM MgCl_2_, 1 mM PMSF) supplemented with 25 U/mL Benzonase nuclease (Sigma-Aldrich, E1014). Lysates were cleared by centrifugation; a fraction of the supernatant was reserved as the input control, and the remaining lysate was incubated with GFP-Trap Magnetic Agarose beads (ChromoTek; gtma-20) for 1 hour at 4 °C with gentle rotation. The beads were washed three times in dilution buffer (50 mM Tris-HCl, pH 7.5, 150 mM NaCl, 0.5 mM EDTA). Bound proteins were eluted and denatured by resuspending the beads in 1× Laemmli sample buffer (Bio-Rad) containing 5% β-mercaptoethanol (Sigma-Aldrich; M6250) and boiling for 5 minutes. The resulting supernatant was analysed by Western blotting.

### Mass spectrometry (MS)

#### Sample preparation and digestion

For the CDC20 immunoprecipitation-mass spectrometry experiment, bead-bound proteins were resuspended in 100 μL of 100 mM triethylammonium bicarbonate (TEAB), reduced by the addition of 1 μL of 0.5 M tris(2-carboxyethyl)phosphine (TCEP), and alkylated with 5 μL of 200 mM iodoacetamide (IAA). Samples were digested with trypsin by adding 10 μL of a 500 ng/μL trypsin solution and incubating overnight with shaking. The supernatant containing the peptides was then transferred to clean microcentrifuge tubes and dried to completeness in a SpeedVac concentrator prior to LC-MS analysis. Nocodazole-arrested RPE1 FRT/TR wild-type and ABBA-mutant PICH-mClover3 cells were collected by mitotic shake-off, lysed, and subjected to GFP pulldown as described in the protocol above. Trypsin digestion and TMT-3plex (Tandem mass tag) mass spectrometry were performed at the Institute of Cancer Research (ICR). In brief, samples were solubilised, reduced, alkylated, and digested, then labelled with TMTpro multiplexing reagents (Thermo Scientific). Prior to LC-MS analysis, TMT-labelled peptides were fractionated by high-pH reversed-phase (RP) chromatography and enriched by immobilised metal ion affinity chromatography. The SequestHT search engine was used to analyse the data.

#### LC-MS/MS data acquisition

GFP-TRAP immunoprecipitates were analysed on an Orbitrap Ascend mass spectrometer (Thermo Fisher Scientific). MS1 spectra were acquired in the Orbitrap at a resolution of 120,000. Precursors were selected for HCD fragmentation in top-speed mode (3 s cycle time) at a normalized collision energy of 32% and detected in the ion trap using the turbo scan rate. MS3 spectra were triggered by real-time search (RTS) against a FASTA database containing reviewed UniProt Homo sapiens canonical and isoform sequences, using multi-notch isolation (10 notches) and HCD fragmentation at 65% normalized collision energy, with detection in the Orbitrap at 120,000 resolution. Dynamic exclusion was enabled for 45 s with a mass tolerance of 10 ppm, RTS close-out was set to a maximum of four peptides per protein, and targeted precursors were excluded from further activation. Static modifications used for RTS included TMTpro16plex on peptide N-termini and lysine residues (+304.2071 Da) and carbamidomethylation of cysteine (+57.0215 Da); variable modifications included oxidation of methionine (+15.9949 Da) and deamidation of asparagine and glutamine (+0.984 Da), with a maximum of one missed cleavage and up to two variable modifications per peptide.

For CDC20 immunoprecipitation, dried peptide samples were resuspended in LC-MS loading buffer and analyzed by nano-liquid chromatography coupled to tandem mass spectrometry using an UltiMate 3000 UHPLC system interfaced with an Orbitrap Fusion Lumos Tribrid mass spectrometer (Thermo Fisher Scientific). Peptides were separated at a flow rate of 300 nL/min over a 120 min gradient, with solvent B (80% acetonitrile, 0.1% formic acid) increasing from 5% to 32% over 95 min, followed by a ramp to 95% B over 5 min, a 5 min wash at 95% B, and re-equilibration at 5% B. The instrument was operated in positive-ion, data-dependent acquisition mode with a 3 s cycle time. Full MS scans were acquired in the Orbitrap at a resolution of 120,000 over an m/z range of 375–1500, with an AGC target of 4 × 10⁵ and a maximum injection time of 50 ms. The most abundant precursor ions with charge states 2–7 were selected for fragmentation by HCD (normalized collision energy 35%) using a 1.2 m/z isolation window. MS/MS spectra were acquired in the ion trap in rapid scan mode, with an AGC target of 2 × 10⁴ and a maximum injection time of 50 ms. Dynamic exclusion was set to 45 s, and precursor ions with intensities below 5,000 were excluded from fragmentation.

#### Database searching and data processing

Raw data were searched using the Sequest HT and Comet nodes in Proteome Discoverer version 3.0 (Thermo Fisher Scientific) against a reviewed UniProt Homo sapiens FASTA database. The precursor mass tolerance was 20 ppm, and the fragment ion mass tolerance was 0.5 Da for Sequest HT and 1 Da for Comet, allowing up to two missed tryptic cleavages. TMTpro modifications on peptide N-termini and lysine residues and carbamidomethylation of cysteine were specified as static modifications, while oxidation of methionine and deamidation of asparagine/glutamine were specified as dynamic modifications. Peptide-spectrum matches were filtered using Percolator at a peptide-level false discovery rate (FDR) of 1% based on a target–decoy strategy. Only unique peptides were used for protein quantification, considering protein groups for peptide uniqueness, and only peptides with an average reporter ion signal-to-noise ratio greater than 3 were included in the quantitative analysis.

For CDC20 pulldown, raw data files were processed in Proteome Discoverer version 2.4 (Thermo Fisher Scientific). MS/MS spectra were searched against a reviewed UniProt human protein sequence database using the Sequest HT search engine with a precursor mass tolerance of 20 ppm and a fragment mass tolerance of 0.5 Da. Trypsin was specified as the proteolytic enzyme, allowing up to two missed cleavages. Carbamidomethylation of cysteine (+57.021 Da) was set as a fixed modification, while oxidation of methionine (+15.995 Da), deamidation of asparagine and glutamine (+0.984 Da), and phosphorylation of serine, threonine, and tyrosine (+79.966 Da) were set as variable modifications. Peptide-spectrum matches were validated using Percolator with a target–decoy strategy, and peptide and protein identifications were filtered to an FDR of 1% at both the peptide and protein levels. Only high-confidence peptide identifications were retained for downstream analyses, and proteins were grouped according to the strict parsimony principle.

### Yeast two-hybrid analysis

For yeast two-hybrid experiments, strain pJ69-4A (ade2-1 his3-200 leu2-3,112 trp1-901 ura3-52 gal4Δ gal80Δ MET2::GAL7-lacZ LYS2::GAL1-HIS3 GAL2-ADE2) was co-transformed with vectors pGBKT7 and pGADT7, each either unmodified or modified to express protein constructs fused to the Gal4-binding or Gal4-activating domain. Logarithmic cultures of the respective strains, grown in minimal medium lacking tryptophan and leucine, were diluted and spotted on control plates (minimal medium lacking tryptophan and leucine) and selective plates (minimal medium lacking tryptophan, leucine, and histidine). The interaction between GAD-TRX-FANCM-MM2 and GBD-RMI2-RMI1(OB3-OB2) and its reciprocal combination were used as a positive control.

### AlphaFold protein structure prediction

For modelling of the PICH-BTRR complex, FANCM-TRR subcomplex and PICH-BLM structures, FASTA protein sequences were submitted to AlphaFold version 2.3.2^62^ run on Apocrita, the Queen Mary, University of London High-Performance Cluster. Except for the PICH-BLM prediction, where 25 models were generated, for the first two complexes, 5 models with 5 seeds were produced using multimer mode, and the top-ranked model was relaxed and taken for figure production using PYMOL Version 3.1.3.1 (The PyMOL Molecular Graphics System, Schrödinger, LLC) and AlphaPickle Version 1.5.4^63^ for statistical plots. The structural modelling of the RMI OB-fold pocket and PICH SLiM, the PICH core domains and BLM RB-2 motif, PICH core domains and double-stranded DNA, and PICH and XPO1 were generated by using AlphaFold3^64^. PAE viewer was used to generate plots of the predicted aligned error^65^.

### ConSurf analysis of Protein Sequence Conservation

ConSurf^66,67^ is run under CoLab to analyse protein sequence conservation using the following parameters: Mode; PDB, Substitution model; choose the best substitution model using protest, rate4site_algorithm; Bayesian. Protein sequence conservation is generated by aligning 150 sequences that represent the list of homologues to the query.

### Metaphase spreads preparation

Cells were first synchronised using a single thymidine block and released into G2/M phase for 6-8 hours. Two hours before the metaphase spread preparation, the cells were treated with 60 nM BI2536, a PLK1 inhibitor (Cayman Chemical; 17385). Mitotic cells were collected by mitotic shake-off. Cell pellets were resuspended and gently mixed in 10 ml of pre-warmed hypotonic solution (0.075 M KCl), then incubated at 37 °C for 10 min. The swollen cells were centrifuged, fixed, and washed twice with Carnoy’s Fixative (3:1 v/v; methanol: acetic acid) according to the standard chromosome spread preparation. Chromosome spreads were dropped onto glass slides and stored at room temperature prior to FISH hybridisation.

### Centromere and Telomere Fluorescence *in Situ* Hybridisation (FISH)

The Peptide nucleic acid (PNA) probe hybridisation of chromosome spreads followed the manufacturer’s guidelines (DAKO Agilent & PNAbio). Chromosome spreads were fixed in 3.7% PFA and washed with a gradient of ice-cold ethanol (70%, 90%, and 100% EtOH). Slides were then air-dried before adding PNA probes to the spreads: centromere, CENTB-FAM (PNAbio; F3006), and Telomere, TelG-Cy3 (PNA FISH kit, Dako Agilent; K532611-8 or PNAbio; F1006). A coverslip (18x18mm) was added to the probe, and the slide was co-denatured at 80 °C for 30 sec to 1 min and hybridised at room temperature in a humidifying chamber for 2 hours. Slides were then washed in Wash Solution (Dako Agilent) at 65°C and dehydrated again in the ice-cold ethanol series. Slides were air-dried and counterstained using Vectashield with DAPI.

### Quantification and statistical analysis

Statistical analysis was performed using GraphPad Prism software version 10.2 by two-tailed unpaired Student’s t-test. Data were presented as the mean ± standard deviation (S.D.) unless specified. The probability value ‘p ≤ 0.05’ is considered to be significant.

## Acknowledgements

We thank Professor Marcel van Vugt (University of Groningen) for the ΔBLM HAP1 cells, Professor Weidong Wang (National Institute on Aging) for the human RMI1 plasmid, Professor Mark Burkard (University of Wisconsin) for the cDNA of CENPB (residue 1-158), and Dr Gary Ying-Wai Chan (the University of Hong Kong) for the CRISPR plasmids for tagging endogenous PICH with mAID-mClover3. We thank Dr Alessandro Bianchi for sharing his expertise in yeast two-hybrid assays. We also acknowledge Dr Zuza Kozik in the Proteomics Core Facility (The Institute of Cancer Research) for the mass spectrometry services. This research utilised Queen Mary’s Apocrita HPC facility, supported by QMUL (Queen Mary University of London) Research-IT. We are thankful to Jonathan Wing for the help with cell sorting (University of Sussex) and to Dr Yan Gu and the Wolfson Centre for Biological Imaging (University of Sussex) for providing excellent microscopy and imaging facilities. We gratefully acknowledge the support of the ICR Core facilities, in particular, light microscopy and flow cytometry. We also thank Dr María Fernández-Casañas for preliminary data analysis and discussion. We thank all members of the Chan and Pines Labs for their valuable ideas and comments on this research. This project was funded by the CRUK Programme grant (C29/A24397) to J.P. and Wellcome Trust Career Development Award (225348/Z/22/Z) to K.-L.C.

## Author contributions

U.A., O.O., M.S., E.K., A.C., Z.K., T.R., A.W.O. and K.-L.C. performed the experiments. U.A., O.O., J.C., A.W.O., J.P. and K.-L.C. designed the experiments. Z.K., T.R. and J.C. performed mass spectrometry analysis. M.D., A.W.O., and K.-L.C. performed and analysed AlphaFold modelling. M.F.C. performed preliminary AlphaFold2 analysis. J.P. and K.-L.C. conceptualised the project and supervised the research. U.A. and K.-L.C. prepared figures and wrote the manuscript with input from all the authors. U.A., O.O., M.D., A.W.O., J.P and K.-L.C. edited the manuscript.

## Data availability

The mass spectrometry proteomics data have been deposited to the ProteomeXchange Consortium via the PRIDE partner repository with the dataset identifier PXD081930.

## Competing interests

The authors declare no competing interests.

---

**Supplementary Figure 1.**
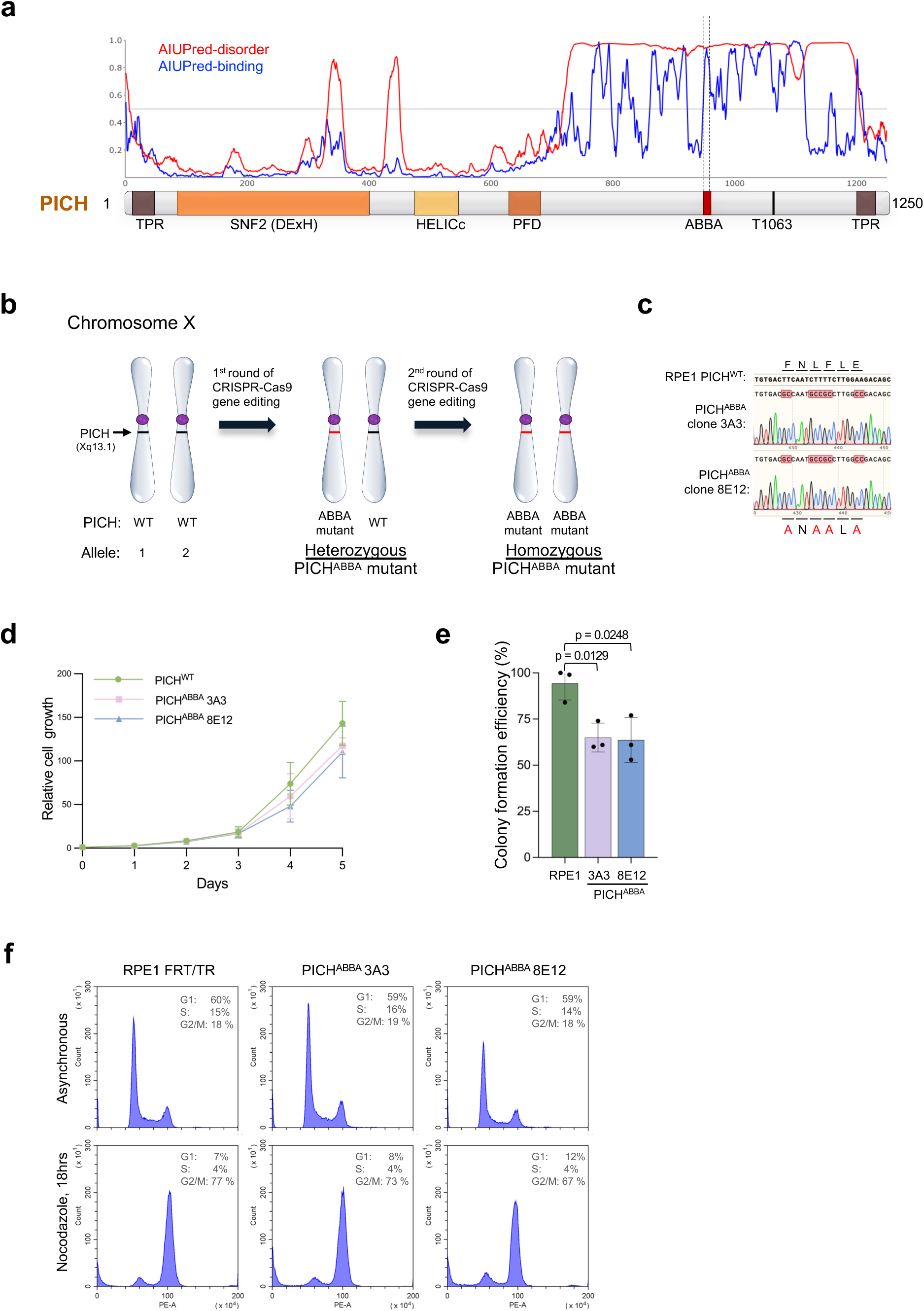
Mutations in the ABBA motif cause a slow-growth phenotype and reduced colony-forming efficiency. **a.** Intrinsically disordered regions and potential protein interaction motifs in PICH predicted by AIUPred. **b.** The CRISPR genome editing approach to create knock-in mutations at both alleles of PICH in RPE1 FRT/TR cells. **c.** Sequencing validation of the CRISPR knock-in mutations of endogenous PICH in two RPE1 mutant clones. **d.** Relative growth rates (to Day 0) of PICH wildtype and the ABBA mutant RPE1 cells. Two mutant clones, 3A3 and 8E12, were generated and analysed. Data presented from two independent biological experiments, each performed in technical duplicates. Mean ± S.D. is shown. **e.** Colony formation efficiencies of PICH wildtype and ABBA mutant RPE1 cells. Data presented from three independent biological experiments, each performed in technical duplicates. Mean ± S.D. is shown. **f.** FACS analysis of PICH wildtype and ABBA mutant RPE1 cells in untreated and nocodazole-treated conditions.

**Supplementary Figure 2.**
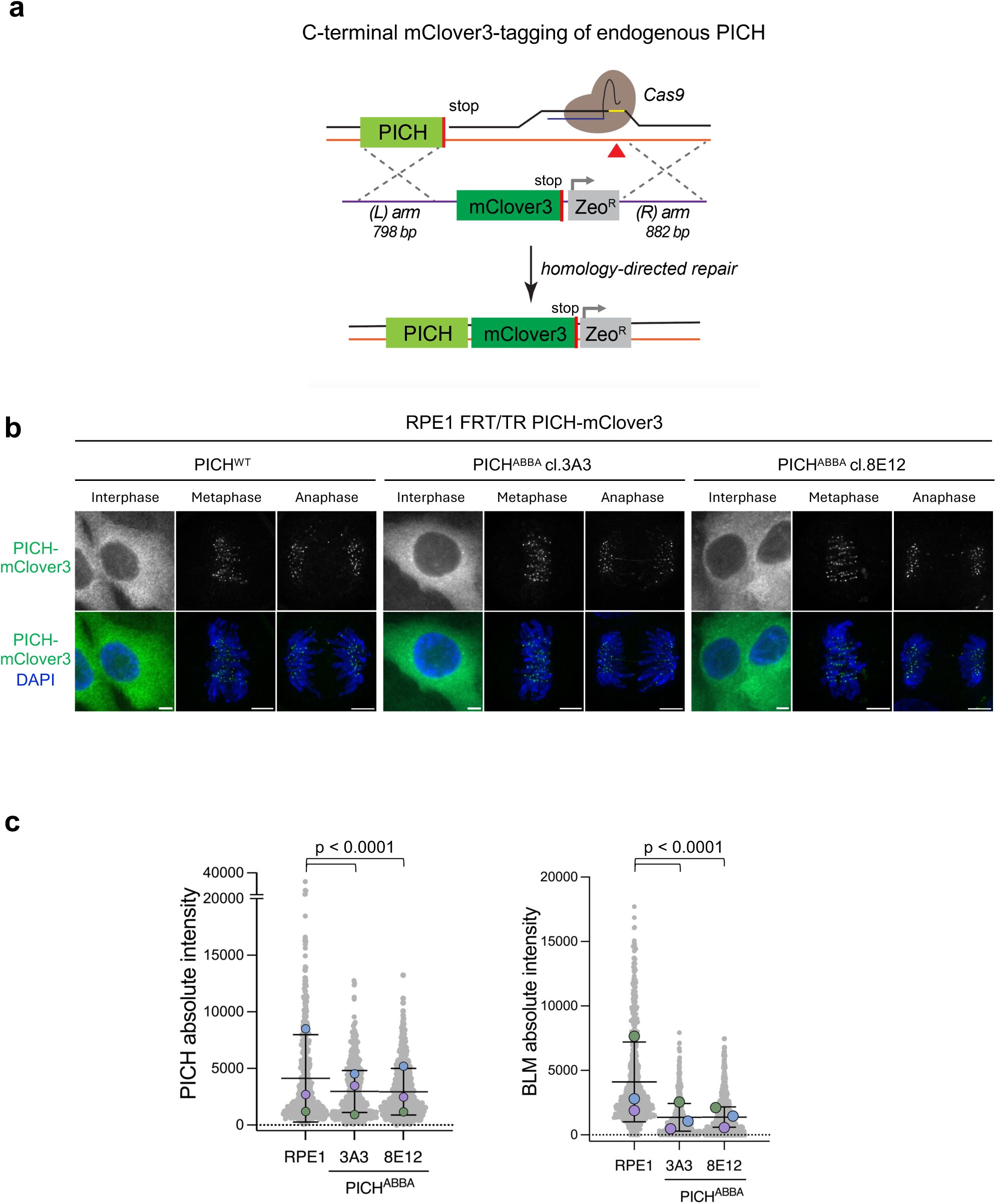
CRISPR-mediated C-terminal mClover3 tagging of endogenous PICH. **a.** A graphical presentation of the strategy of mClover3-tagging at the C-terminus of endogenous PICH in wildtype and ABBA mutant RPE1 cells by CRISPR technology. **b.** Representative images of the localisation of mClover3-tagged endogenous PICH in interphase and mitotic cells. **c.** Measurements of PICH and BLM intensities on UFBs in PICH^WT^ and PICH^ABBA^ RPE1 cells (see **Fig.1d**).

**Supplementary Figure 3.**
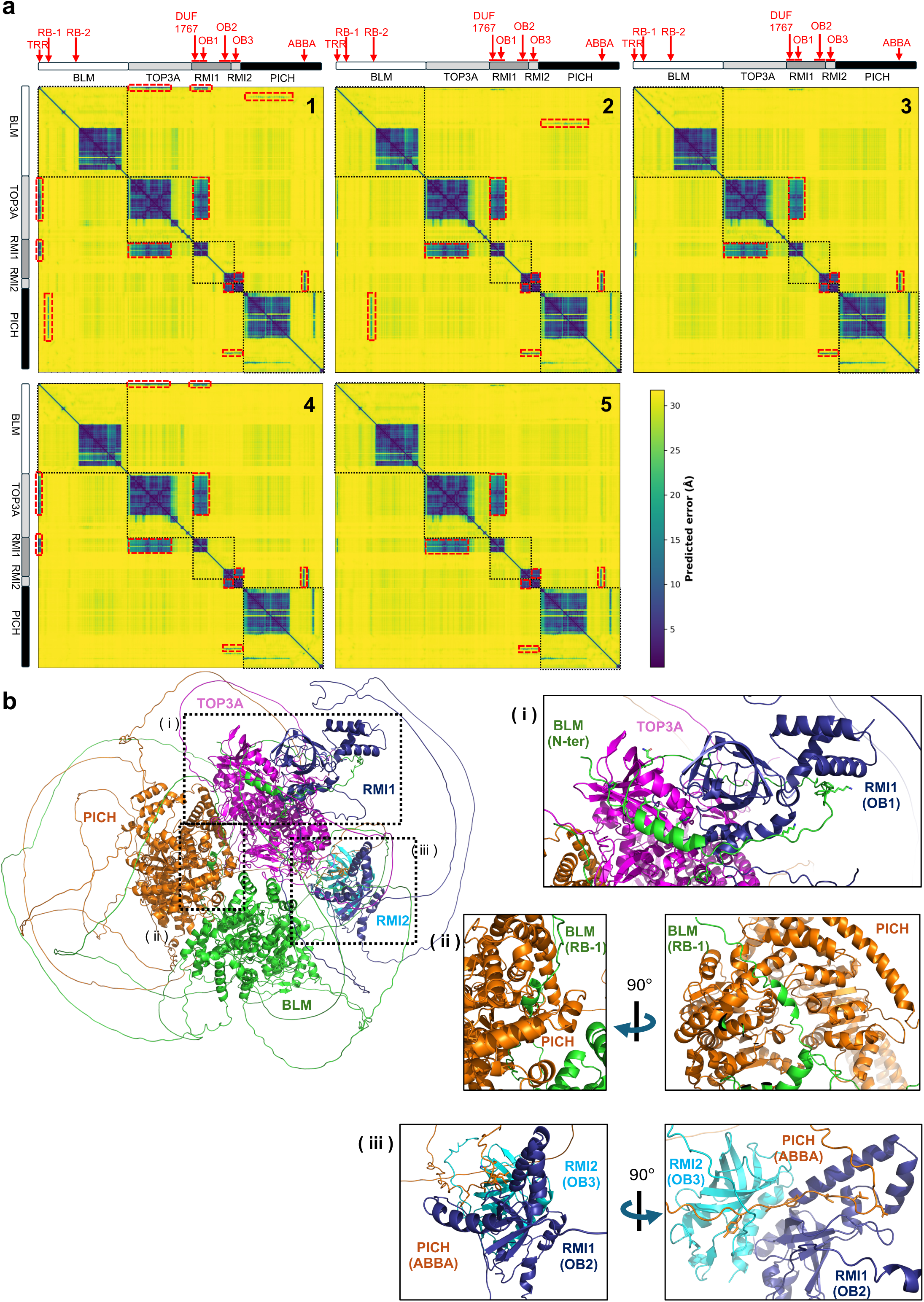
AlphaFold-Multimer modelling of the UFB-binding complex components. **a.** Predicted Aligned Errors (PAEs) of the AF-Multimer models of the UFB-binding factors including BLM, TOP3A, RMI1, RMI2 and PICH. Five models are shown. Red boxes indicate potential intermolecular protein-protein interactions between the indicated domains or regions of the subunits. **b.** An example of the structural model made from prediction 1. Dotted boxes showing the interactions (i) of the N-terminus of BLM with TOP3A and RMI1 OB-fold domain 1 (OB-1), (ii) between the BLM RPA-binding motif 1 (RB-1) and the core domains of PICH, and (iii) between the PICH ABBA motif and the heterodimer of RMI1 OB-fold domain 2 (OB-2) and RMI2 OB-fold domain 3 (OB-3).

**Supplementary Figure 4.**
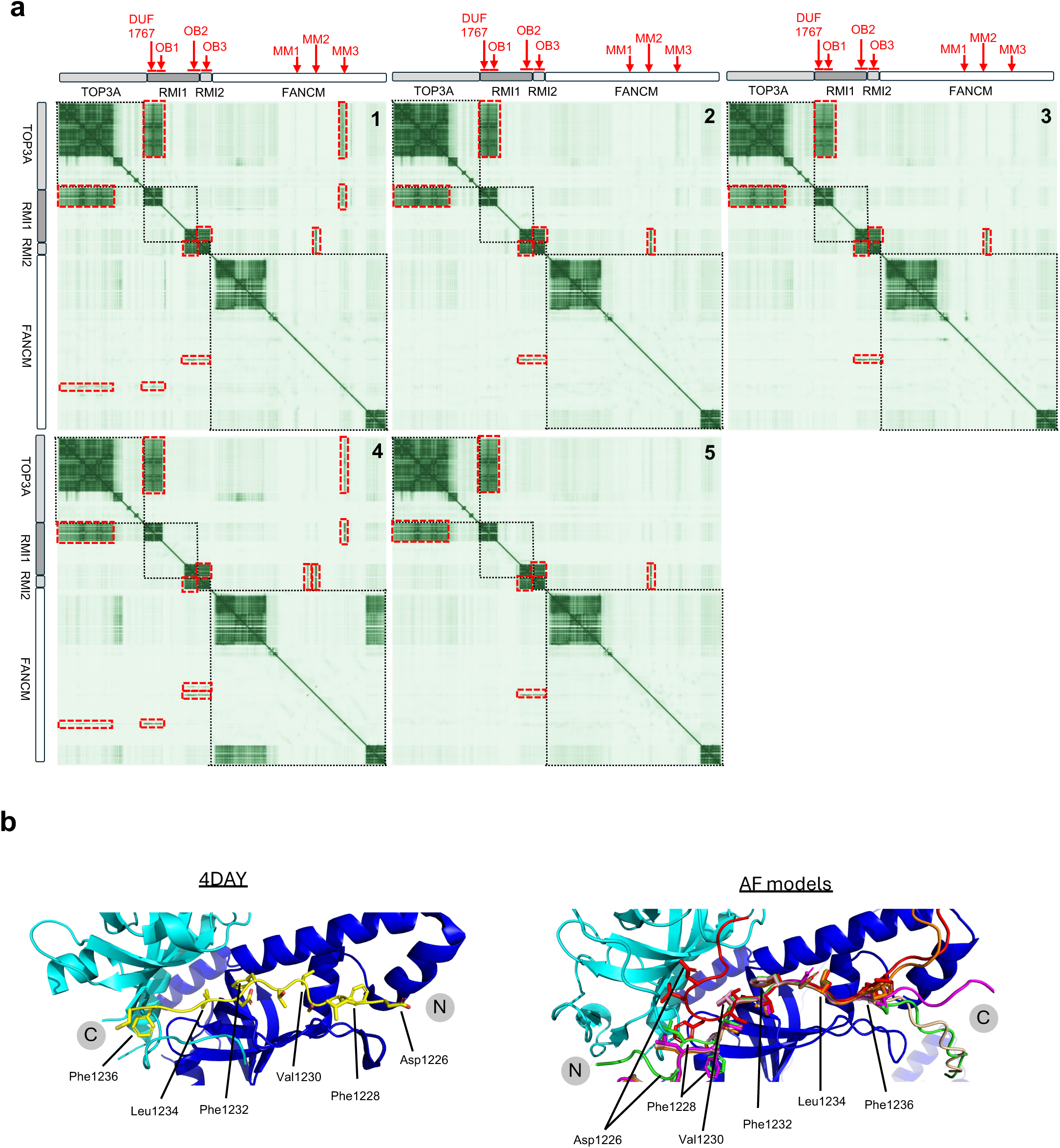
AlphaFold-Multimer modelling of the interaction among TOP3A, RMI1, RMI2 and FANCM. **a.** Predicted Aligned Errors (PAEs) of the five AF-Multimer models. Red boxes indicate potential intermolecular protein-protein interactions between the indicated domains or regions of the subunits. **b.** Opposite binding orientations of the FANCM MM2 motif to the RMI OB-fold pocket in the co-crystal (4DAY) and AF-Multimer predicted structures. Conserved residues responsible for the RMI OB-fold pocket binding are shown.

**Supplementary Figure 5.**
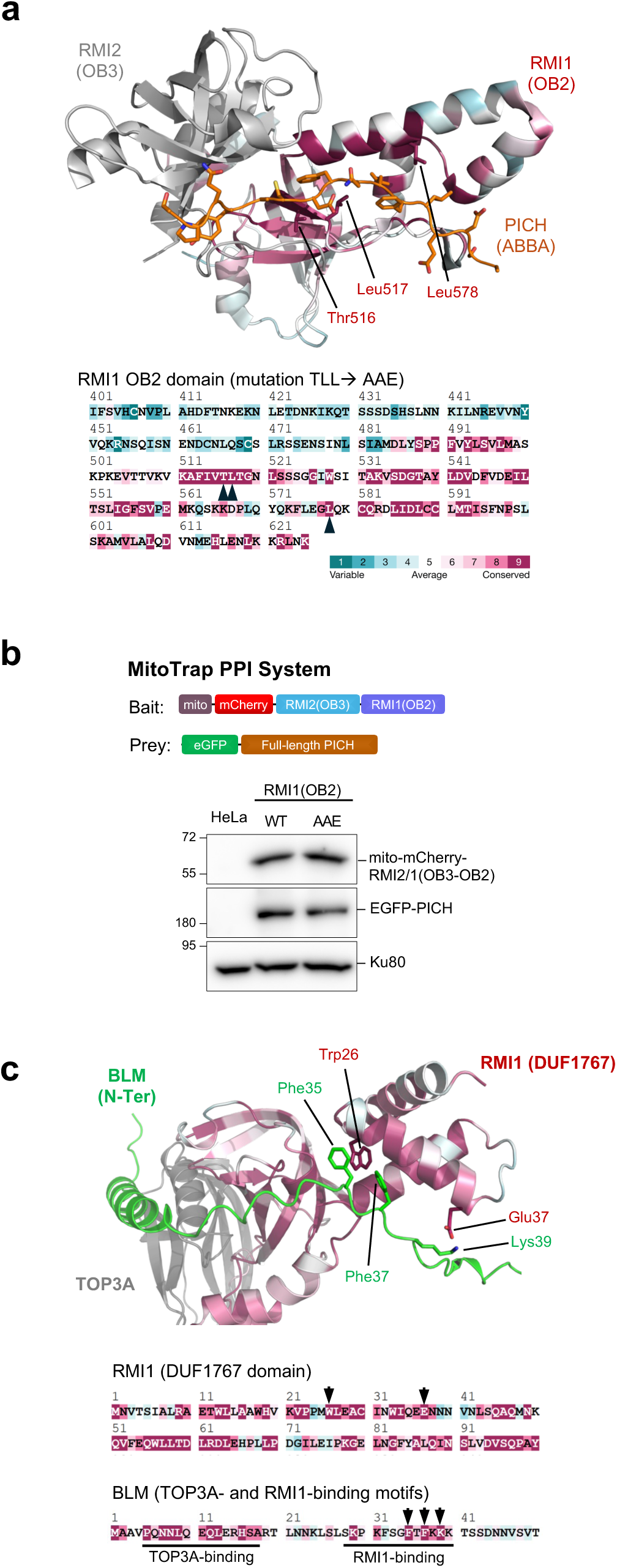
AlphaFold3 and ConSurf prediction of conserved residues in RMI1 for the interaction with PICH SLiM and BLM. **a.** ConSurf analysis reveals the conserved residues within the AlphaFold3 model of the RMI1 OB2 domain in a complex with RMI2 for PICH ABBA motif interaction. Arrowheads: residues (Thr516, Leu517 and Leu578) were mutated for protein-protein interaction analysis. **b.** MitoTrap PPI analysis between the RMI OB-fold pocket (fused to mito-mCherry) and full-length PICH (fused to eGFP). Western blots showing expression of the fusion proteins in HeLa cells. Ku80 is used as a loading control. **c.** AlphaFold3 modelling of the interaction of the N-terminus of BLM with TOP3A and RMI1 (DUF1767) domain, respectively. Trp26 and Glu37 of RMI1 are predicted to be essential for the interaction with Phe35, Phe37 and Lys39 in BLM. ConSurf analysis showing the conserved residues in the RMI1(DUF1767) domain and the N-terminus of BLM. Arrows indicate the potential residues responsible for the RMI1-BLM interaction. Trp26 and Glu37 of RMI1 are chosen for mutagenesis to inactive BLM binding.

**Supplementary Figure 6.**
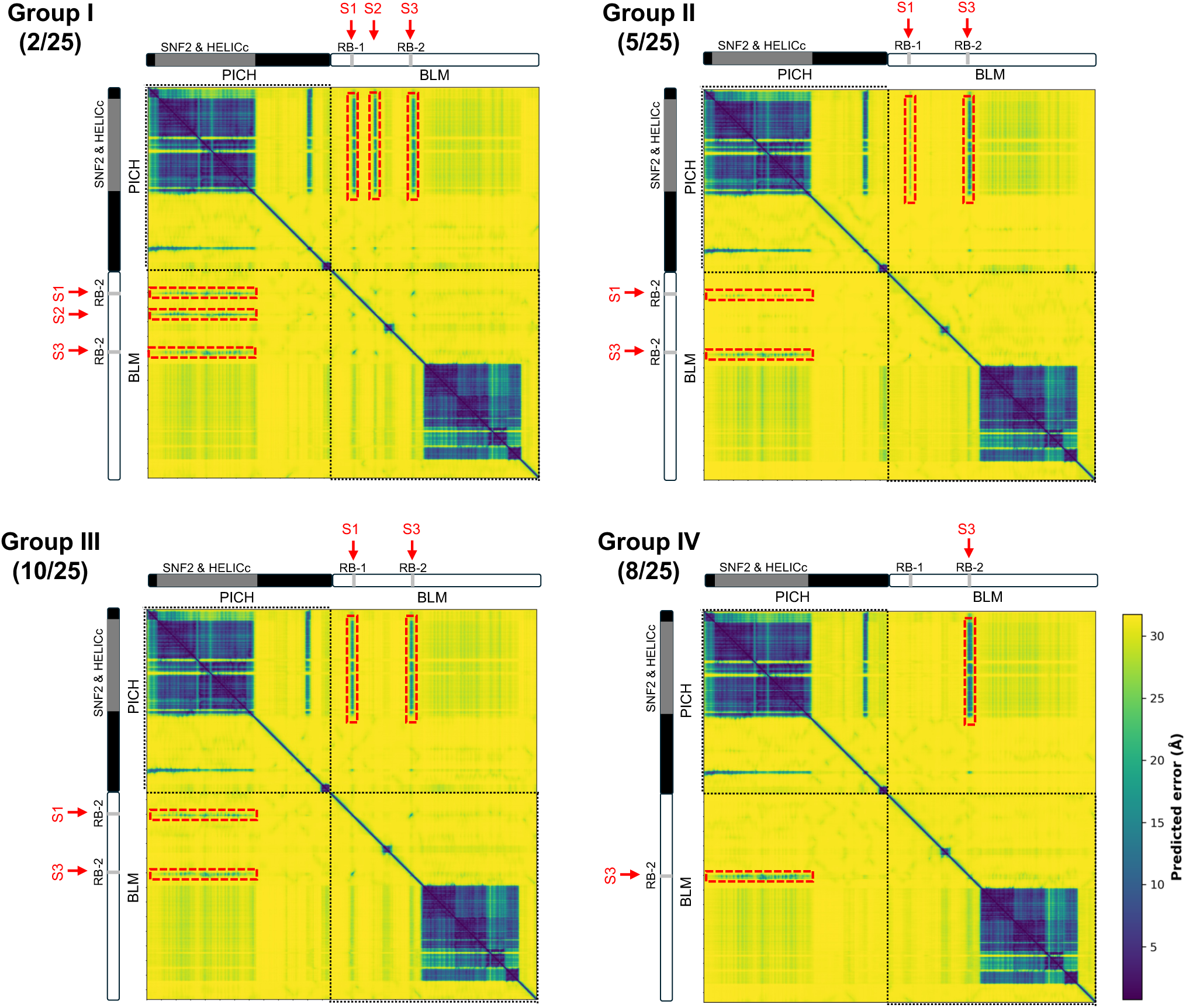
AlphaFold-Multimer modelling of the interaction between PICH and BLM. Predicted Aligned Errors of the twenty-five AF-Multimer models. They are classified into 4 groups based on the number and level of potential protein-protein interactions. Red boxes indicate the potential sites (S1-3) in BLM, which show potential interactions with the core domains of PICH.

**Supplementary Figure 7.**
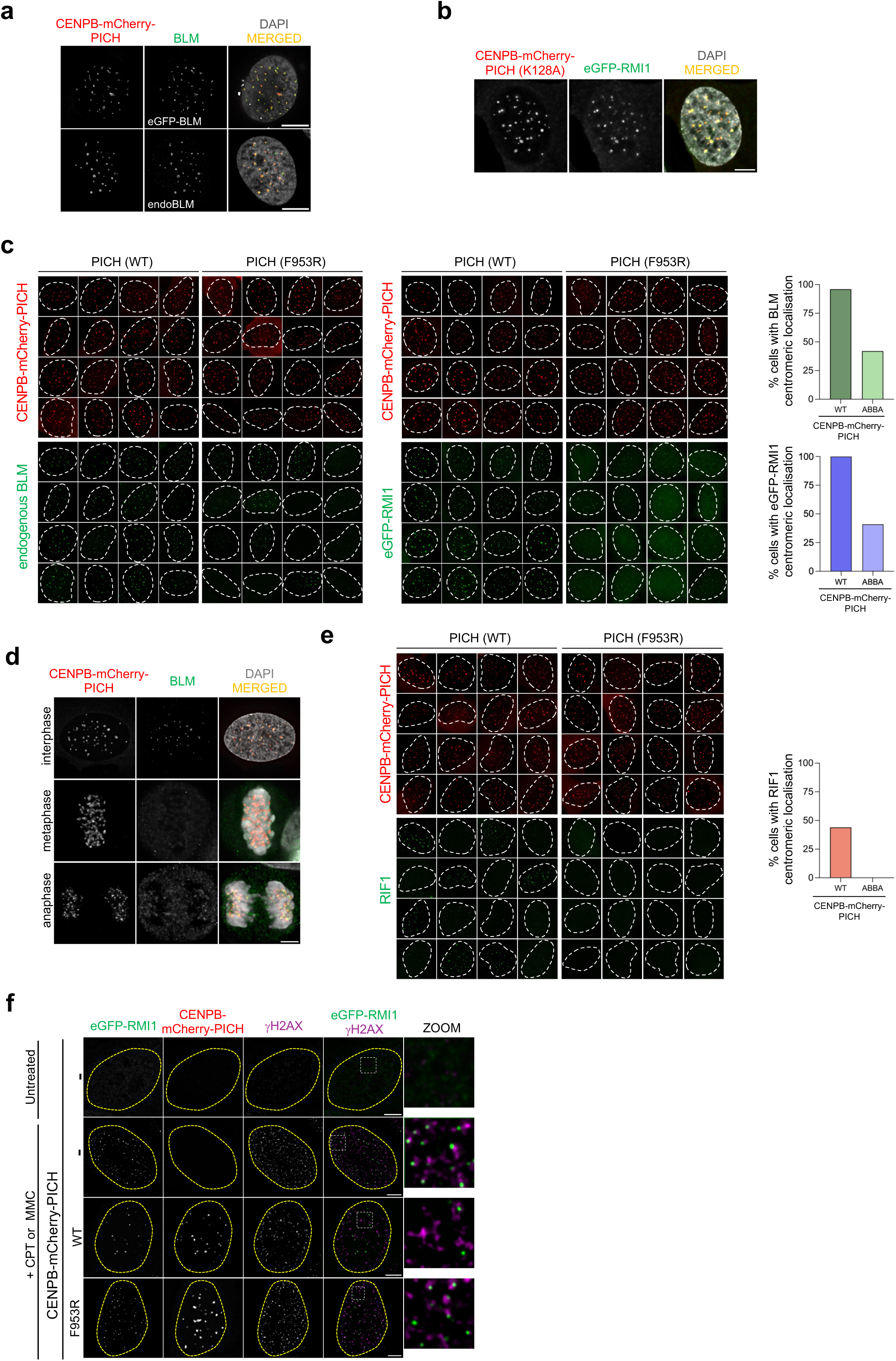
**a.** Immunofluorescent staining images showing the recruitment of eGFP-tagged and endogenous BLM to centromeres by CENPB-mCherry-PICH in HeLa cells. **b.** Immunofluorescent staining images showing recruitment of eGFP-tagged RMI1 to centromeres by an active-site dead mutant of PICH(K128A) fused with CENPB-mCherry in HeLa cells. **c.** Gallery of representative images showing recruitment of eGFP-tagged RMI1 and endogenous BLM to centromeres by CENPB-mCherry-PICH^WT^ or PICH^F953R^ in HeLa cells. Right: percentage of cells showing eGFP-RMI1 and BLM recruitment to centromeres. Total number (n) of cells analysed: eGFP-RMI1 PICH^WT^, n = 117; eGFP-RMI1 PICH^F953R^, n = 74; endogenous BLM PICH^WT^, n = 107; endogenous BLM PICH^F953R^, n = 126. **d.** Immunofluorescent staining images showing recruitment of endogenous BLM to centromeres by CENPB-mCherry-PICH in interphase and mitotic HeLa cells. **e.** Gallery of representative images showing recruitment of endogenous RIF1 to centromeres by CENPB-mCherry-PICH^WT^ or PICH^F953R^ in HeLa cells. Right: percentage of cells showing RIF1 recruitment to centromeres. Total number (n) of cells analysed: PICH^WT^, n = 126; PICH^F953R^, n = 118. **f.** Immunofluorescence images showing HeLa cells with or without the expression of CENPB-mCherry-PICH^WT^ or PICH^F953R^ and the localisation of eGFP-RMI1 at DNA damage sites (marked by γH2AX) after the indicated treatments. Scale bars, 10 μm.

**Supplementary Figure 8.**
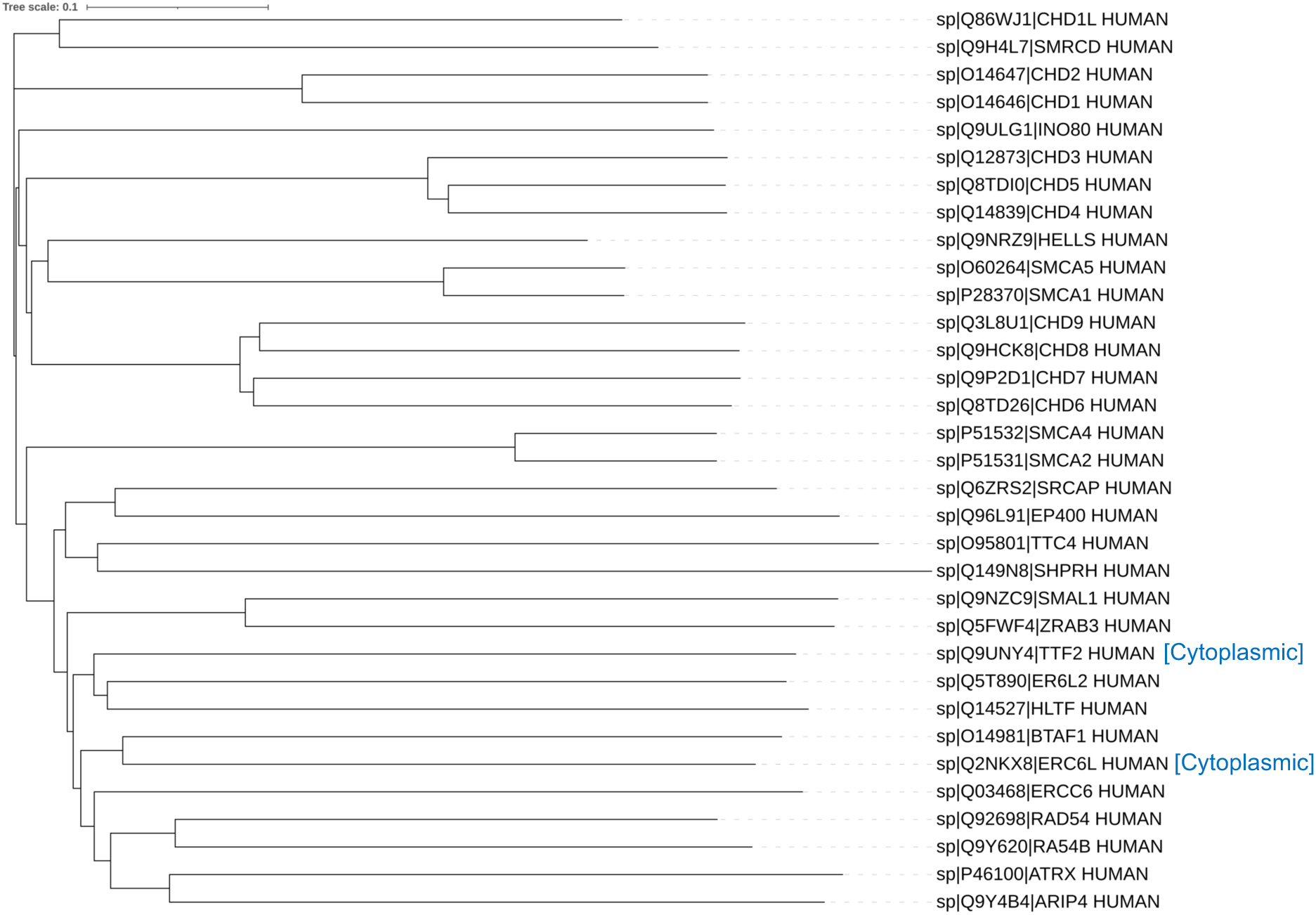
Phylogenetic tree of all known human SNF2-like DNA translocases generated by MUSCLE/iTOL software.

## Notes

### Competing Interest Statement

The authors have declared no competing interest.

## References

1 Bythell-Douglas, R. & Deans, A. J. A Structural Guide to the Bloom Syndrome Complex. Structure 29, 99–113, doi:10.1016/j.str.2020.11.020 (2021).

2 Chan, K. L., North, P. S. & Hickson, I. D. BLM is required for faithful chromosome segregation and its localization defines a class of ultrafine anaphase bridges. EMBO J 26, 3397–3409, doi:10.1038/sj.emboj.7601777 (2007).

3 Fernandez-Casanas, M. & Chan, K. L. The Unresolved Problem of DNA Bridging. Genes (Basel) 9, doi:10.3390/genes9120623 (2018).

4 Hamann, T. E. et al. Mitotic BLM functions are required to maintain genomic stability. Nucleic Acids Res 54, doi:10.1093/nar/gkag199 (2026).

5 Shorrocks, A. K. et al. The Bloom syndrome complex senses RPA-coated single-stranded DNA to restart stalled replication forks. Nat Commun 12, 585, doi:10.1038/s41467-020-20818-5 (2021).

6 Wu, L. & Hickson, I. D. The Bloom’s syndrome helicase suppresses crossing over during homologous recombination. Nature 426, 870–874, doi:10.1038/nature02253 (2003).

7 Chaudhury, I., Sareen, A., Raghunandan, M. & Sobeck, A. FANCD2 regulates BLM complex functions independently of FANCI to promote replication fork recovery. Nucleic Acids Res 41, 6444–6459, doi:10.1093/nar/gkt348 (2013).

8 Patel, D. S., Misenko, S. M., Her, J. & Bunting, S. F. BLM helicase regulates DNA repair by counteracting RAD51 loading at DNA double-strand break sites. J Cell Biol 216, 3521–3534, doi:10.1083/jcb.201703144 (2017).

9 Bussen, W., Raynard, S., Busygina, V., Singh, A. K. & Sung, P. Holliday junction processing activity of the BLM-Topo IIIalpha-BLAP75 complex. J Biol Chem 282, 31484–31492, doi:10.1074/jbc.M706116200 (2007).

10 Ho, H. N. & West, S. C. Generation of double Holliday junction DNAs and their dissolution/resolution within a chromatin context. Proc Natl Acad Sci U S A 119, e2123420119, doi:10.1073/pnas.2123420119 (2022).

11 Chan, K. L., Palmai-Pallag, T., Ying, S. & Hickson, I. D. Replication stress induces sister-chromatid bridging at fragile site loci in mitosis. Nat Cell Biol 11, 753–760, doi:10.1038/ncb1882 (2009).

12 Naim, V. & Rosselli, F. The FANC pathway and BLM collaborate during mitosis to prevent micro-nucleation and chromosome abnormalities. Nat Cell Biol 11, 761–768, doi:10.1038/ncb1883 (2009).

13 Ellis, N. A. et al. The Bloom’s syndrome gene product is homologous to RecQ helicases. Cell 83, 655–666, doi:10.1016/0092-8674(95)90105-1 (1995).

14 Ellis, N. A. & German, J. Molecular genetics of Bloom’s syndrome. Hum Mol Genet 5 Spec No, 1457–1463, doi:10.1093/hmg/5.supplement_1.1457 (1996).

15 Martin, C. A. et al. Mutations in TOP3A Cause a Bloom Syndrome-like Disorder. Am J Hum Genet 103, 221–231, doi:10.1016/j.ajhg.2018.07.001 (2018).

16 Bergeron, K. L., Murphy, E. L., Brown, L. W. & Almeida, K. H. Critical interaction domains between bloom syndrome protein and RAD51. Protein J 30, 1–8, doi:10.1007/s10930-010-9295-8 (2011).

17 Srivastava, V. et al. BLM helicase stimulates the ATPase and chromatin-remodeling activities of RAD54. J Cell Sci 122, 3093–3103, doi:10.1242/jcs.051813 (2009).

18 Blackford, A. N. et al. TopBP1 interacts with BLM to maintain genome stability but is dispensable for preventing BLM degradation. Mol Cell 57, 1133–1141, doi:10.1016/j.molcel.2015.02.012 (2015).

19 Tripathi, V., Kaur, S. & Sengupta, S. Phosphorylation-dependent interactions of BLM and 53BP1 are required for their anti-recombinogenic roles during homologous recombination. Carcinogenesis 29, 52–61, doi:10.1093/carcin/bgm238 (2008).

20 Nimonkar, A. V. et al. BLM-DNA2-RPA-MRN and EXO1-BLM-RPA-MRN constitute two DNA end resection machineries for human DNA break repair. Genes Dev 25, 350–362, doi:10.1101/gad.2003811 (2011).

21 Nimonkar, A. V., Ozsoy, A. Z., Genschel, J., Modrich, P. & Kowalczykowski, S. C. Human exonuclease 1 and BLM helicase interact to resect DNA and initiate DNA repair. Proc Natl Acad Sci U S A 105, 16906–16911, doi:10.1073/pnas.0809380105 (2008).

22 Deans, A. J. & West, S. C. FANCM connects the genome instability disorders Bloom’s Syndrome and Fanconi Anemia. Mol Cell 36, 943–953, doi:10.1016/j.molcel.2009.12.006 (2009).

23 Hoadley, K. A. et al. Defining the molecular interface that connects the Fanconi anemia protein FANCM to the Bloom syndrome dissolvasome. Proc Natl Acad Sci U S A 109, 4437–4442, doi:10.1073/pnas.1117279109 (2012).

24 Baumann, C., Korner, R., Hofmann, K. & Nigg, E. A. PICH, a centromere-associated SNF2 family ATPase, is regulated by Plk1 and required for the spindle checkpoint. Cell 128, 101–114, doi:10.1016/j.cell.2006.11.041 (2007).

25 Hengeveld, R. C. et al. Rif1 Is Required for Resolution of Ultrafine DNA Bridges in Anaphase to Ensure Genomic Stability. Dev Cell 34, 466–474, doi:10.1016/j.devcel.2015.06.014 (2015).

26 Addis Jones, O., Tiwari, A., Olukoga, T., Herbert, A. & Chan, K. L. PLK1 facilitates chromosome biorientation by suppressing centromere disintegration driven by BLM-mediated unwinding and spindle pulling. Nat Commun 10, 2861, doi:10.1038/s41467-019-10938-y (2019).

27 Fernandez-Casanas, M. et al. Centromere protection requires strict mitotic inactivation of the Bloom syndrome helicase complex. Nat Commun 16, 7832, doi:10.1038/s41467-025-62966-6 (2025).

28 Lera, R. F. et al. Plk1 protects kinetochore-centromere architecture against microtubule pulling forces. EMBO Rep 20, e48711, doi:10.15252/embr.201948711 (2019).

29 Biebricher, A. et al. PICH: a DNA translocase specially adapted for processing anaphase bridge DNA. Mol Cell 51, 691–701, doi:10.1016/j.molcel.2013.07.016 (2013).

30 Spakman, D. et al. PICH acts as a force-dependent nucleosome remodeler. Nat Commun 13, 7277, doi:10.1038/s41467-022-35040-8 (2022).

31 Ke, Y. et al. PICH and BLM limit histone association with anaphase centromeric DNA threads and promote their resolution. EMBO J 30, 3309–3321, doi:10.1038/emboj.2011.226 (2011).

32 Eustermann, S., Patel, A. B., Hopfner, K. P., He, Y. & Korber, P. Energy-driven genome regulation by ATP-dependent chromatin remodellers. Nat Rev Mol Cell Biol 25, 309–332, doi:10.1038/s41580-023-00683-y (2024).

33 Erdos, G. & Dosztanyi, Z. AIUPred: combining energy estimation with deep learning for the enhanced prediction of protein disorder. Nucleic Acids Res 52, W176–W181, doi:10.1093/nar/gkae385 (2024).

34 Di Fiore, B. et al. The ABBA motif binds APC/C activators and is shared by APC/C substrates and regulators. Dev Cell 32, 358–372, doi:10.1016/j.devcel.2015.01.003 (2015).

35 Di Fiore, B., Wurzenberger, C., Davey, N. E. & Pines, J. The Mitotic Checkpoint Complex Requires an Evolutionary Conserved Cassette to Bind and Inhibit Active APC/C. Mol Cell 64, 1144–1153, doi:10.1016/j.molcel.2016.11.006 (2016).

36 Hubner, N. C. et al. Re-examination of siRNA specificity questions role of PICH and Tao1 in the spindle checkpoint and identifies Mad2 as a sensitive target for small RNAs. Chromosoma 119, 149–165, doi:10.1007/s00412-009-0244-2 (2010).

37 Kong, N. et al. The interplay of the translocase activity and protein recruitment function of PICH in ultrafine anaphase bridge resolution and genomic stability. Nucleic Acids Res 53, doi:10.1093/nar/gkae1249 (2025).

38 Robinson, M. S., Sahlender, D. A. & Foster, S. D. Rapid inactivation of proteins by rapamycin-induced rerouting to mitochondria. Dev Cell 18, 324–331, doi:10.1016/j.devcel.2009.12.015 (2010).

39 Evans, R., et al. Protein complex prediction with AlphaFold-Multimer. bioRxiv, 2021.2010.2004.463034, doi:10.1101/2021.10.04.463034 (2022).

40 Raynard, S. et al. Functional role of BLAP75 in BLM-topoisomerase IIIalpha-dependent holliday junction processing. J Biol Chem 283, 15701–15708, doi:10.1074/jbc.M802127200 (2008).

41 Wu, L. et al. The Bloom’s syndrome gene product interacts with topoisomerase III. J Biol Chem 275, 9636–9644, doi:10.1074/jbc.275.13.9636 (2000).

42 Wang, F. et al. Crystal structures of RMI1 and RMI2, two OB-fold regulatory subunits of the BLM complex. Structure 18, 1159–1170, doi:10.1016/j.str.2010.06.008 (2010).

43 Hoadley, K. A. et al. Structure and cellular roles of the RMI core complex from the bloom syndrome dissolvasome. Structure 18, 1149–1158, doi:10.1016/j.str.2010.06.009 (2010).

44 Yang, J., O’Donnell, L., Durocher, D. & Brown, G. W. RMI1 promotes DNA replication fork progression and recovery from replication fork stress. Mol Cell Biol 32, 3054–3064, doi:10.1128/MCB.00255-12 (2012).

45 Sarlos, K. et al. Reconstitution of anaphase DNA bridge recognition and disjunction. Nat Struct Mol Biol 25, 868–876, doi:10.1038/s41594-018-0123-8 (2018).

46 Hodson, C. et al. Mechanism of Bloom syndrome complex assembly required for double Holliday junction dissolution and genome stability. Proc Natl Acad Sci U S A 119, doi:10.1073/pnas.2109093119 (2022).

47 Guo, R. B., Rigolet, P., Zargarian, L., Fermandjian, S. & Xi, X. G. Structural and functional characterizations reveal the importance of a zinc binding domain in Bloom’s syndrome helicase. Nucleic Acids Res 33, 3109–3124, doi:10.1093/nar/gki619 (2005).

48 Kong, N., Liu, Z. & Chan, Y. W. RIF1 suppresses the formation of single-stranded ultrafine anaphase bridges via protein phosphatase 1. Cell Rep 42, 112032, doi:10.1016/j.celrep.2023.112032 (2023).

49 Yoda, K., Kitagawa, K., Masumoto, H., Muro, Y. & Okazaki, T. A human centromere protein, CENP-B, has a DNA binding domain containing four potential alpha helices at the NH2 terminus, which is separable from dimerizing activity. J Cell Biol 119, 1413–1427, doi:10.1083/jcb.119.6.1413 (1992).

50 Kosugi, S. et al. Six classes of nuclear localization signals specific to different binding grooves of importin alpha. J Biol Chem 284, 478–485, doi:10.1074/jbc.M807017200 (2009).

51 Kosugi, S., Hasebe, M., Tomita, M. & Yanagawa, H. Systematic identification of cell cycle-dependent yeast nucleocytoplasmic shuttling proteins by prediction of composite motifs. Proc Natl Acad Sci U S A 106, 10171–10176, doi:10.1073/pnas.0900604106 (2009).

52 Dong, X. et al. Structural basis for leucine-rich nuclear export signal recognition by CRM1. Nature 458, 1136–1141, doi:10.1038/nature07975 (2009).

53 Davey, N. E. et al. Attributes of short linear motifs. Mol Biosyst 8, 268–281, doi:10.1039/c1mb05231d (2012).

54 Karow, J. K., Newman, R. H., Freemont, P. S. & Hickson, I. D. Oligomeric ring structure of the Bloom’s syndrome helicase. Curr Biol 9, 597–600, doi:10.1016/s0960-9822(99)80264-4 (1999).

55 Shi, J. et al. A helical bundle in the N-terminal domain of the BLM helicase mediates dimer and potentially hexamer formation. J Biol Chem 292, 5909–5920, doi:10.1074/jbc.M116.761510 (2017).

56 Tian, T. et al. The ZATT-TOP2A-PICH Axis Drives Extensive Replication Fork Reversal to Promote Genome Stability. Mol Cell 81, 198–211 e196, doi:10.1016/j.molcel.2020.11.007 (2021).

57 Kee, Y., Kim, J. M. & D’Andrea, A. D. Regulated degradation of FANCM in the Fanconi anemia pathway during mitosis. Genes Dev 23, 555–560, doi:10.1101/gad.1761309 (2009).

58 Lu, R. et al. The FANCM-BLM-TOP3A-RMI complex suppresses alternative lengthening of telomeres (ALT). Nat Commun 10, 2252, doi:10.1038/s41467-019-10180-6 (2019).

59 Alcock, L. J. et al. Potent Cyclic Peptide Inhibitors Disrupt the FANCM-RMI Interaction. J Med Chem 68, 12615–12625, doi:10.1021/acs.jmedchem.5c00365 (2025).

60 Alcock, L. J. et al. Cell-Active Peptide Inhibitors of the FANCM-RMI Interaction. J Med Chem 68, 24672–24683, doi:10.1021/acs.jmedchem.5c02655 (2025).

61 Chiang, T. W., le Sage, C., Larrieu, D., Demir, M. & Jackson, S. P. CRISPR-Cas9(D10A) nickase-based genotypic and phenotypic screening to enhance genome editing. Sci Rep 6, 24356, doi:10.1038/srep24356 (2016).

62 Jumper, J. et al. Highly accurate protein structure prediction with AlphaFold. Nature 596, 583–589, doi:10.1038/s41586-021-03819-2 (2021).

63 Arnold, M. J. AlphaPickle (2021).

64 Abramson, J. et al. Accurate structure prediction of biomolecular interactions with AlphaFold 3. Nature 630, 493–500, doi:10.1038/s41586-024-07487-w (2024).

65 Elfmann, C. & Stulke, J. PAE viewer: a webserver for the interactive visualization of the predicted aligned error for multimer structure predictions and crosslinks. Nucleic Acids Res 51, W404–W410, doi:10.1093/nar/gkad350 (2023).

66 Ashkenazy, H. et al. ConSurf 2016: an improved methodology to estimate and visualize evolutionary conservation in macromolecules. Nucleic Acids Res 44, W344–350, doi:10.1093/nar/gkw408 (2016).

67 Landau, M. et al. ConSurf 2005: the projection of evolutionary conservation scores of residues on protein structures. Nucleic Acids Res 33, W299–302, doi:10.1093/nar/gki370 (2005).

